# Targeting Cblb-Notch1 axis as a novel immunotherapeutic strategy to boost CD8+ T-cells responses

**DOI:** 10.1101/2022.03.15.484378

**Authors:** Giulia Monticone, Fred Csibi, Silvana Leit, David Ciccone, Ameya S. Champhekar, Jermaine E. Austin, Deniz A. Ucar, Fokhrul Hossain, Zhi Huang, Salome V. Ibba, A. Hamid Boulares, Nicholas Carpino, Samarpan Majumder, Keli Xu, Barbara A. Osborne, Christine Loh, Lucio Miele

**Affiliations:** Louisiana State University Health Sciences Center, Department of Genetics, New Orleans, LA, United States; Nimbus Therapeutics, Cambridge, MA, United States; University of California, Los Angeles, CA, United States; University of Virginia, Charlottesville, VA, United States; Louisiana State University Health Sciences Center, Department of Interdisciplinary Oncology, New Orleans, LA, United States; Stony Brook University, Stony Brook, NY, United States; University of Mississippi Medical Center, Department of Neurobiology and Anatomical Sciences, Jackson, MS, United States; University of Massachusetts, Department of Veterinary and Animal Sciences, Amherst, MA, United States

**Keywords:** Adenosine, Cbl-b, Immunotherapy, Immunosuppression, Notch1

## Abstract

A critical feature of cancer is the ability to induce immunosuppression and evade immune responses. Tumor-induced immunosuppression diminishes the efficacy of endogenous immune responses and decreases the efficacy of cancer immunotherapy. In this study, we describe a new immunosuppressive pathway in which adenosine promotes Cbl-b-mediated Notch1 degradation, causing suppression of CD8+ T-cells effector functions. Genetic KO and pharmacological inhibition of Cbl-b prevents Notch1 degradation in response to adenosine and reactivates its signaling. Reactivation of Notch1 results in enhanced CD8+ T-cell effector functions, anti-cancer response and resistance to immunosuppression. Our work demonstrates that targeting Cbl-b-Notch1 axis is a novel promising strategy for cancer immunotherapy.

## Introduction

Tumor-induced immunosuppression is a hallmark feature of cancer which allows tumors to evade immune surveillance and progress (Hanahan, 2011). This is a major challenge for endogenous anti-cancer immune responses as well as for the successful application of cancer immunotherapies (Hegde, 2020). Therefore, there is a critical need for novel cancer immunotherapies that can not only boost the immune response, but also overcome tumor-induced immunosuppression.

Tumors achieve immunosuppression in different ways, including production of immunosuppressive molecules, recruitment of suppressive immune cells and formation of physical barriers to immune infiltration (Labani-Motlagh, 2020). Among these strategies, overproduction of adenosine, an ATP metabolite, plays a major part in suppressing immune responses in the tumor microenvironment (Vijayan, 2017). Adenosine modulates the immune response by activating transmembrane adenosine receptors expressed on the membrane of immune cells (Otha, 2001; Linden, 2006). Several studies have shown that adenosine suppresses CD8+ T-cells by activating the adenosine A2A receptor (A2AR) and blocking A2AR with selective antagonists results in enhanced anti-cancer immune responses (Sorrentino, 2019; Willingham, 2018; Vijayan, 2017; Ohta, 2006). Recent evidence has shown that A2AR activation leads to downregulation of Notch1, a key regulator of T-cell effector functions (Sorrentino, 2019). Notch1 signaling is triggered by T-cell receptor (TCR) activation and it modulates T-cell effector functions by regulating proliferation and production of cytokines, including γ-interferon (INF-γ) and Granzyme B (GNZB) (Steinbuck, 2018; Dongre, 2014; Cho, 2009; Palaga, 2003). In line with the positive role of Notch1 in regulating T-cell functions, exogenous expression of Notch1 in CD8+ T-cells enhances anti-cancer T-cells responses and render T-cells resistant to immunosuppression in the tumor microenvironment of lung carcinoma and thymoma (Sierra, 2014). On the contrary, inhibition of Notch1 activation with gamma-secretase inhibitors (GSI) has been shown to suppress T-cell activation, reduce proliferation and cytokine production (Majumder, 2021; Golde, 2013; Kopan, 2004). These findings suggest that Notch1 is a potential target to control tumor-induced immunosuppression.

Notch receptors are transmembrane proteins consisting of a large extracellular domain for ligand binding, a transmembrane domain and an intracellular domain that exerts transcriptional regulation. Upon binding to ligands presented on adjacent cells, Notch goes through proteolytic cleavages mediated by ADAM10 and gamma-secretase, respectively, releasing Notch intracellular domain (NICD) which will translocate into the cell nucleus, form a complex with co-activators and activate transcription of target genes (Bray, 2016; Chillakuri, 2012). Notch signaling can be also regulated in a ligand-independent manner (Monticone, 2021; Palmer, 2015; Baron, 2012). Ligand-independent endocytic regulation of Notch is of particular importance in *Drosophila* (Shimizu, 2014; Vaccari, 2009; Wilkin, 2004), as well as, in mammalian T-cells, where TCR activation triggers ligand-independent activation of Notch1 (Steinbuck, 2018; Cho, 2009; Palaga, 2003). Several studies have reported that different ubiquitin ligases are involved in the ligand-independent regulation of Notch in *Drosophila* (Shimizu, 2014; Dalton, 2011; Wilkin, 2004). However, it is yet unknown which and how ubiquitin ligases are involved in regulating Notch1 in T-cells.

Casitas B-lineage lymphoma b (Cbl-b) is an E3 ubiquitin ligase that has been identified as an important negative regulator of TCR signaling cascade. Cbl-b controls the threshold of T-cell activation and T-cell anergy (Paolino, 2011; Chiang, 2000). Cbl-b deficient T-cells have a lower activation threshold and can be stimulated in the absence of CD28 co-stimulation (Chiang, 2000). Mice lacking Cbl-b reject tumors because of increased T-cells and NK cells immune response (Paolino, 2014; Chiang, 2007; Loeser, 2007) Depletion of Cbl-b inhibits exhaustion in CD8+ T-cells and CAR-T cells (Kumar, 2021).

Here, we describe a new regulatory axis in which adenosine, via A2AR, promotes Cbl-b-mediated Notch1 degradation, causing immunosuppression in CD8+ T-cells. Genetic KO and pharmacological inhibition of Cbl-b prevents Notch1 degradation in response to adenosine and reactivates its signaling. Notch1 reactivation results in enhanced CD8+ T-cell effector functions, anti-cancer response and resistance to immunosuppression. Our findings indicate that A2AR-Cbl-b-Notch1 axis is a novel promising target for cancer immunotherapy.

## Results

### A2AR antagonism promotes Notch1-positive CD8+ T-cells anti-tumor responses

Recent studies have proposed that A2AR signaling negatively regulates Notch1 in CD8+ T-cells (Sorrentino, 2019). Other studies have shown that A2AR blockade with selective antagonists in T-cells results in enhanced anti-tumor activity (Willingham, 2018; Vijayan, 2017; Ohta, 2006). In light of these findings, we hypothesized that A2AR antagonists promote anti-tumor activity in CD8+ T cells through restoring Notch1 function. To test this idea, we generated tumor-derived organoids from a syngeneic Triple-Negative-Breast Cancer (TNBC) model, C0321 (Zhang, 2014; Zhang, 2015). Tumor-derived organoids are clusters of cells which contain all cell types present in the original tumor, including cancer cells, infiltrating immune cells and stroma cells. Several studies have shown that tumor-derived organoids are a reliable *ex-vivo* system that recapitulates the features of the original tumor and its microenvironment (Jenkins, 2019; Puca, 2018). In this study, TNBC tumor-derived organoids were treated with a selective A2AR antagonist, ZM-241395 (ZM), and a selective A2AR agonist CGS-21680 (CGS) to mimic physiological adenosine. A2AR blockade by ZM significantly suppressed tumor organoid growth compared to control and CGS-treated organoids (Fig. 1A). In addition, ZM was effective only in organoids derived from immunocompetent mice, but not from immunocompromised athymic Nu/Nu mice that have a vastly reduced number of T-cells, suggesting that the anti-tumor effect of ZM is dependent on T-cells (Fig. 1B). We marked CD8+ T-cells in organoids with an anti-CD8 fluorophore-conjugated antibody and found that ZM-treated organoids displayed higher infiltration and clusters of CD8+ T-cells surrounding cancer cells, instead of the dispersed organization of T-cells observed in control organoids (Fig. 1C, Supplementary fig. 1). This pattern of CD8+ T-cells might reflect cytotoxic responses in ZM-treated organoids. Remarkably, more CD8+ T-cells were positive for Notch1 in ZM-treated organoids than in controls (Fig. 1C), suggesting that ZM promotes CD8+ T-cells infiltration and activation in a Notch1-dependent manner. Overall, these results indicate that blocking A2AR signaling with selective antagonists leads to enhanced anti-tumor activity of Notch1-positive CD8+ T-cells.

**Fig. 1.**
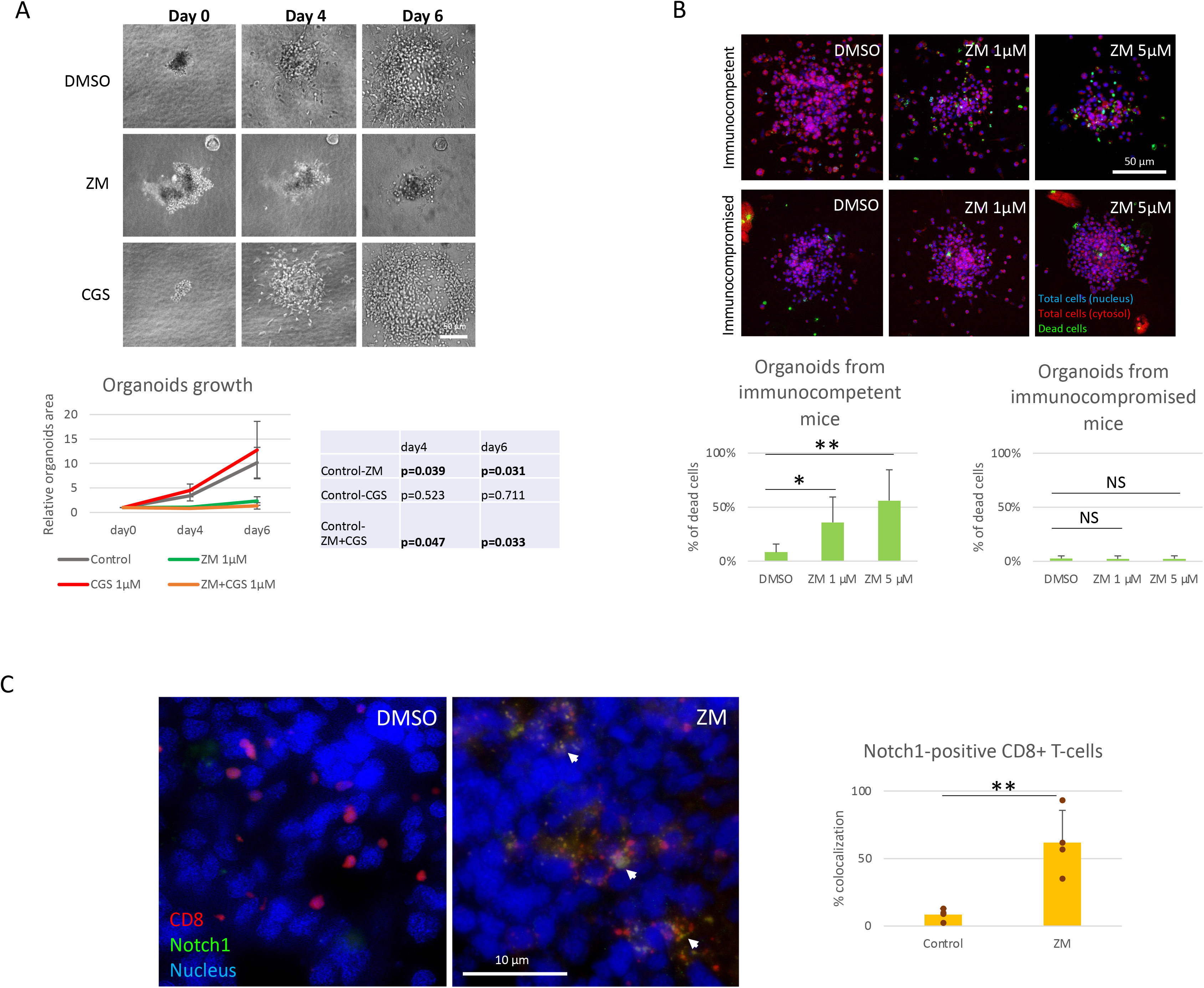
A2AR antagonist ZM enhances Notch1-positive T-cells anti-tumor activity. (A) Organoids were derived from a syngeneic TNBC mouse model, C0321, in FVB mice, cultured in a collagen matrix for 6 days and treated with vehicle (DMSO) or 1µM ZM-241385 (ZM), 1µM CGS-21680 (CGS) or ZM+CGS. The growth of organoids was determined over time by measuring the area of the same organoids at day 0, 4 and 6. (B) Organoids were obtained from C0321 tumors from immunocompetent FVB mice or immunocompromised athymic nude mice, cultured and treated for 4 days with vehicle, 1µM or 5µM ZM. On day 4, organoids were stained with Hoechst 33342 (nucleus), CellTracker Red (cytosol), NucGreen Dead 488 (dead cells) and imaged. The percentage of dead cells was measured using BZ-x800 analyzer software. (C) C0321 organoids were fixed, permeabilized and stained for CD8+ T-cells and Notch1 with primary anti-CD8, anti-Notch1 and secondary antibodies, and DAPI (nucleus). The colocalization of Notch1 and CD8 was measured using ImageJ software. All images were acquired on a BZ-x800 microscope with a 20x (A,B) and 60x (C) objectives. Scale bars length is indicated above each bar (µm). The graphs show averages ± standard deviation from ≥10 organoids from independent experiments. *p<0.05, **p<0.01,***p<0.001, two tailed T-test with equal variance. NS, non-significant.

### A2AR modulates Notch1 degradation and T-cell functions in CD8+ T-cells

To confirm the previous finding that A2AR activation by adenosine downregulates Notch1 in CD8+ T cells, primary CD8+ T-cells were isolated from mouse spleens and activated with anti-CD3/CD28. To mimic the effect of adenosine, we treated the cells with the selective A2AR agonist CGS. We observed that Notch1 full length (N1FL) and membrane-bound (N1TM) were downregulated in CD8+ T-cells treated with CGS, whereas the A2AR antagonist ZM rescued CGS-mediated downregulation (Fig. 2A). In our previous study, we did not observe effects of CGS on Notch1 mRNA (Sorrentino, 2019), suggesting that Notch1 downregulation may occur at the protein level. To test whether Notch1 downregulation by CGS was the result of increased protein degradation, Notch1 was immunoprecipitated and ubiquitination was detected. We observed that CGS dramatically increased the ubiquitination of Notch1, compared to untreated and ZM-treated CD8+ T-cells (Fig. 2B). To confirm that increased ubiquitination was related to more degradation, CD8+ T-cells were treated with protein synthesis inhibitor Cycloheximide (CHX) and Notch1 was detected at different time points. We found that Notch1 was decreased more rapidly in CGS treated CD8+ T-cells (Fig. 2C), further suggesting that A2AR activation promotes Notch1 degradation. Finally, we tested if A2AR activation results in immunosuppression. We found that CGS decreased proliferation and INF-gamma production in CD8+ T-cells (Fig. 2D). In agreement with the data presented in Fig. 1, these effects can be reversed by ZM (Fig. 2D), indicating that restoring Notch1 drives activation of T-cell function including cell cycle progression and cytokine secretion. Taken together, A2AR activation suppresses CD8+ T-cells function, at least in part, through promoting Notch1 degradation.

**Fig. 2.**
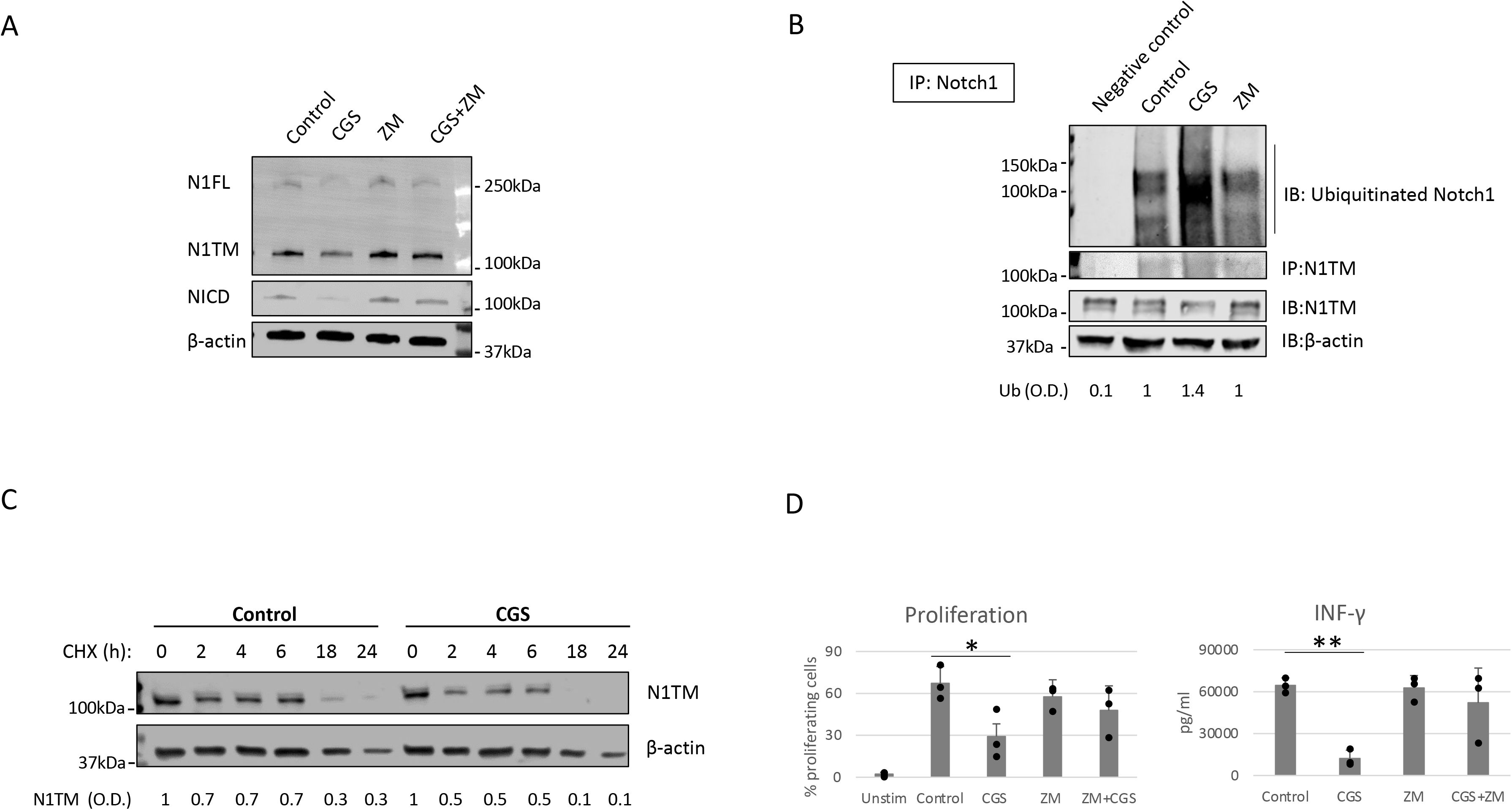
A2AR modulates Notch1 degradation and T-cell functions. (A) Primary CD8+ T-cells isolated from the spleen and lymph nodes of C57B16 or FVB mice were stimulated with anti-CD3/CD28 antibodies and treated with vehicle (control, DMSO) or 1µM ZM-241385 (ZM), 1µM CGS-21680 (CGS) or ZM+CGS for 72h. Protein lysates were analyzed for Notch1 full length (N1FL), Notch1 transmembrane (N1TM) and Notch1 transcriptionally active form (NICD). (B) Primary isolated CD8+ T-cells were stimulated with anti-CD3/CD28 antibodies and treated with vehicle (control, DMSO), 1µM CGS, ZM 1µM for 48h. Notch1 was immunoprecipitated from protein lysates to detect ubiquitinated Notch1. From top to bottom the panels show: ubiquitinated Notch1, N1TM from IP samples, and N1TM, β-actin from input samples. Densitometric (O.D.) results for ubiquitinated Notch1 normalized by IP:N1TM are shown below the panel. Negative control refers to samples immunoprecipitated using beads but not antibody. (C) Primary isolated CD8+ T-cells were stimulated with anti-CD3/CD28 antibodies in the presence of the protein synthesis inhibitor Cycloheximide (CHX) and treated with vehicle (control, DMSO) or 1µM CGS for 24h. N1TM was detected in protein lysates extracted at different time points of CHX treatment, as indicated. Densitometric (O.D.) results for N1TM normalized by β-actin are shown below the panel. (D) Proliferation was measure with flow cytometry by labelling primary CD8+ T-cells with CFSE in samples treated as in (A). Production of INF-γ was analyzed using ELISA in supernatants from samples treated as described in (A). The graphs show averages ± standard deviation from three independent experiments. *p<0.05, **p<0.01, two tailed T-test with equal variance. Unstimulated cells (Unstim) and vehicle treated cells (control) were used as controls. β-actin was used to normalize densitometric values.

### Cbl-b mediates Notch1 degradation in CD8+ T-cells

Having found that A2AR modulates the ubiquitination and degradation of Notch1, we next asked what ubiquitin ligase is involved in regulating Notch1 in CD8+ T-cells. We analyzed the interactome of Notch1 in activated primary CD8+ T-cells using mass spectrometry (Fig. 3A) and found that Cbl-b, a key negative regulator of T-cell functions (Paolino, 2011; Chiang, 2000) was among the ubiquitin ligases that interact the most with Notch1. Indeed, we also found that Cbl-b protein level was increased in CD8+ T-cells treated with CGS and it negatively correlated with Notch1 levels (Supplementary fig. 2).

**Fig. 3.**
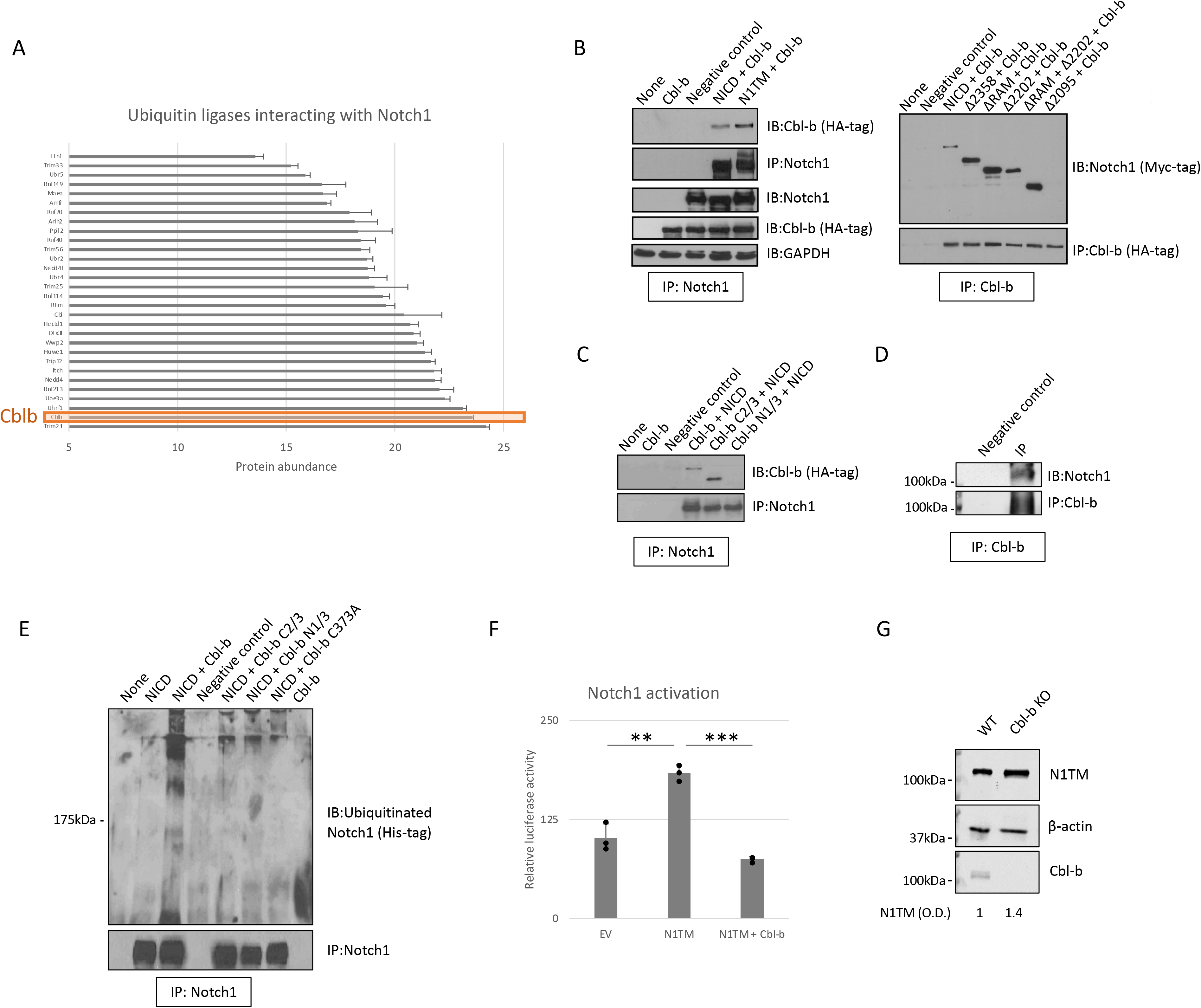
Cbl-b ubiquitinates and degrades Notch1. (A) Mass spectrometry analysis of Notch1 interactome showing ubiquitin ligases interacting with Notch1 and their rate of interaction (protein abundance). The analysis was carried out in lysates from primary CD8+ T-cells stimulated with anti-CD3/CD28 antibodies for 48h, in which Notch1 was immunoprecipitated. Protein lysates were analyzed for Notch1 full length (N1FL) and Notch1 transmembrane (N1TM). (B-C) 293T cells were transfected with various Myc-tagged Notch1 and/or HA-tagged Cbl-b constructs. Lysates were prepared after 48 hours and subjected to immunoprecipitation of Notch1 or Cbl-b, as indicated. (B) In the left panel, Notch1 was immunoprecipitated using a specific anti-Notch1 antibody and Cbl-b was detected with anti-HA tag antibody. From top to bottom the panels show: Cbl-b, Notch1 from IP samples, and Notch1, Cbl-b, GAPDH from input samples. In the right panel, Cbl-b was immunoprecipitated using anti-HA-tag antibody and Notch1 was detected with anti-Myc antibody. The top panel shows Notch1 and the bottom Cbl-b in IP samples. (C) Notch1 was immunoprecipitated using anti-Notch1 antibody and Cbl-b was detected using anti-HA-tag antibody. The top panel shows Cbl-b and the bottom Notch1 in IP samples. (D) Cbl-b was immunoprecipitated and Notch1 detected in lysates from primary isolated CD8+ T-cells stimulated with anti-CD3/CD28 antibodies for 48h. (E) 293T cells were transfected with various Myc-tagged Notch1 and/or HA-tagged Cbl-b constructs as in (B-C) and a His-tagged ubiquitin construct. Notch1 was immunoprecipitated and its ubiquitination was detected with an anti-His tag antibody. The top panel shows ubiquitinated Notch1 and the bottom Notch1 in IP samples. (F) 293T cells were transfected with N1TM with or without Cbl-b construct, as shown. All cells received the Hes-luciferase reporter gene construct. Lysates were prepared after 48 hours and luciferase activity was measured using a luminometer. Empty vector (EV) refers to cells transfected with an empty vector as a negative control. (G) Notch1 was detected in protein lysates from primary CD8+ T-cells isolated from C57BL6 (WT) or Cbl-b -/- (Cbl-b KO) mice and activated with anti-CD3/CD28 antibodies for 72h. From top to bottom the panels show: N1TM, β-actin and Cbl-b. Densitometric (O.D.) results for N1TM normalized by β-actin are shown below the panel. The graphs show averages ± standard deviation from three independent experiments. **p<0.01, ***p<0.001 two tailed T-test with equal variance. Negative control refers to samples immunoprecipitated using rabbit IgG instead of anti-Notch1/HA-tag/Cbl-b antibody. None, non-transfected cells.

To confirm the interaction between Notch1 and Cbl-b and map the interacting sites, we set up a series of immunoprecipitation assays using 293T cells transfected with various Myc-tagged Notch1 and HA-tagged Cbl-b constructs (Fig. 3B, C). We tested whether Cbl-b could associate with membrane bound Notch1 (N1TM) as well as the cleaved, transcriptionally active form of Notch1 (NICD). Cbl-b was pulled down with both Notch constructs, however, N1TM always showed slightly higher affinity for Cbl-b than NICD at similar expression levels (Fig. 3B). Next, we attempted to identify the regions indispensable for Notch1 - Cbl-b interaction. First, 293T cells were transfected with a series of Notch1 constructs in which an increasing amount of the C-terminal region was deleted, along with the HA-tagged full length Cbl-b construct. Among the various deletion mutants used, only Δ2095, a construct that has part of the transcriptional activation domain (TAD) deleted, did not bind Cbl-b (Fig. 3B). Reciprocal experiments using HA-tagged Cbl-b deletion mutants revealed that the C-terminal region of Cbl-b is important for binding Notch1 (Fig. 3C). Deletion of the protein kinase binding domain (TKB) at the N-terminus of Cbl-b in mutant Cbl-b C2/3, did not impair the binding with Notch1, whereas a mutant consisting of TKB only (Cbl-b N1/3) was unable to pull down Notch1 (Fig. 3C). Lastly, Cbl-b could be immunoprecipitated with Notch1 in primary CD8+ T-cells (Fig. 3D). These data showed that Cbl-b and Notch1 associate with each other and a region containing part of the TAD domain of Notch1 and the C-terminal region of Cbl-b are important for this interaction.

As Notch1 ubiquitination is increased in CD8+ T-cells in response to A2AR activation, we asked if Cbl-b regulates Notch1 signaling by directly ubiquitinating Notch1 protein. To do this, 293T cells were transfected with a His-tagged ubiquitin construct along with Notch1 and Cbl-b constructs, as indicated (Fig. 3E). We detected the presence of ubiquitinated Notch1 only in the cells that were co-transfected with Notch1 and full-length Cbl-b. N- and C-terminal deletion mutants, as well as a point mutant of Cbl-b (C373A) which is deficient in its E3 ligase activity, were unable to ubiquitinate Notch1 (Fig. 3E).

To determine whether Cbl-b-mediated Notch1 ubiquitination leads to Notch1 signaling downregulation, we used a reporter gene assay based on a reporter construct in which the luciferase gene is placed downstream of a target of Notch1, the *Hes* promoter (Jarriault, 1998). Reporter gene activity was reduced two- to three-fold when Notch1 and Cbl-b were co-transfected in 293T cells, compared to Notch1 alone (Fig. 3F), indicating that Cbl-b negatively regulated Notch1 signaling. In agreement with these results, we found that Notch1 is upregulated in Cbl-b KO primary CD8+ T-cells compared to WT cells (Fig. 3G), suggesting that lack of Cbl-b rescues Notch1 from degradation. Taken together, these results indicate that Cbl-b ubiquitinates and degrades Notch1 and lack of Cbl-b is sufficient to restore Notch1 protein levels.

### A2AR coordinates Cbl-b-mediated Notch1 degradation via STS-1

To determine how Cbl-b-mediated Notch1 degradation is regulated by A2AR, we analyzed the protein interactome of Notch1 in activated primary CD8+ T-cells treated with CGS or ZM (Fig. 4A, supplementary Table 1). From the mass spectrometry analysis, Suppressor of T-cell receptor signaling 1 (STS-1 or UBASH3B) appeared as the Notch1-interacting protein that was changed the most by the treatments and more importantly, in opposite directions: CGS increased Notch1 interaction with STS-1, whereas ZM decreased this interaction (Fig. 4A,B). This finding was of particular interest because STS-1 is a Tyr-phosphatase that acts as a negative regulator of T-cell activation and is known to interact with Cbl-family proteins through its SH3 domain (Mikhailik, 2007; Carpino, 2004; Kowanetz, 2004). Since Cbl-b activity is negatively regulated by Tyr-phosphorylation (Xiao, 2015), we hypothesized that STS-1 may promote Cbl-b activity by reducing its Tyr-phosphorylation in response to A2AR activation. To test this idea, we immunoprecipitated Cbl-b and detected its Tyr-phosphorylation in response to A2AR activation by CGS. We found that Cbl-b was less Tyr-phosphorylated in CGS-treated CD8+ T-cells compared to untreated cells (Fig. 4C), suggesting that A2AR regulates Cbl-b activity via STS-1-mediated Tyr-dephosphorylation. To confirm that STS-1 acts as a molecular switch that controls Notch1 degradation in response to A2AR activation, we detected Notch1 in STS-1/2 KO CD8+ T-cells treated with CGS. In line with our hypothesis, we found that Notch1 was not downregulated in STS-1/2 KO T-cells in response to CGS treatment (Fig. 4D), indicating that lack of STS-1 prevents A2AR-mediated Notch1 degradation. Our data supports a model in which A2AR controls Cbl-b-mediated Notch1 degradation via STS-1 dephosphorylation and enhancement of Cbl-b activity.

**Fig. 4.**
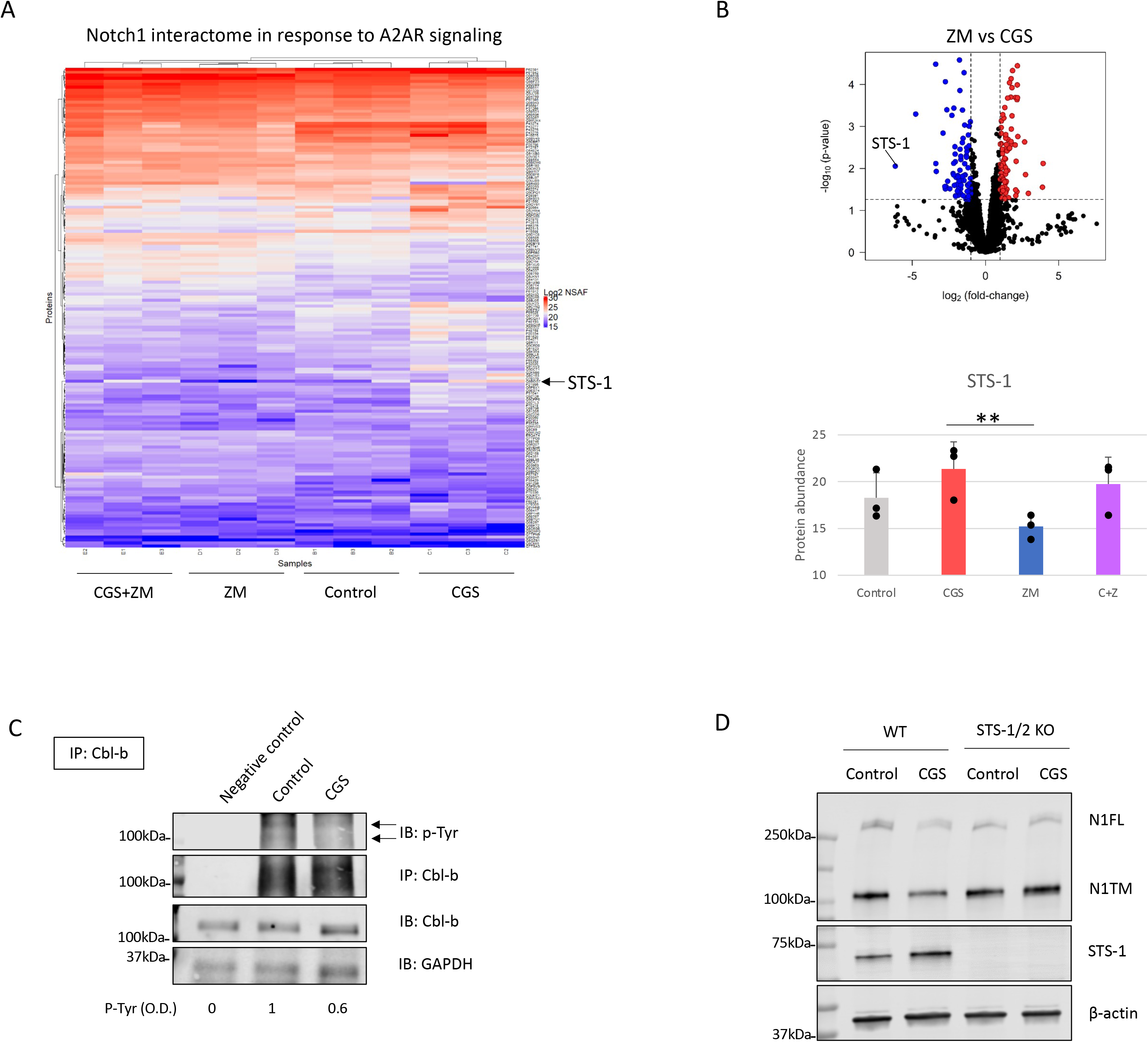
A2AR coordinates Cbl-b-mediated Notch1 degradation via STS-1. The analysis was carried out in lysates from primary mouse CD8+ T-cells stimulated with anti-CD3/CD28 and treated with vehicle (control, DMSO) or 1µM ZM-241385 (ZM), 1µM CGS-21680 (CGS) or ZM+CGS for 48h. Notch1 was immunoprecipitated in the lysates before mass spectrometry analysis. Three independent replicates were analyzed for each sample. (A) Heat map of mass spectrometry analysis of Notch1 interactome. Uniprot IDs of the detected proteins are indicated on the left side of the heat map. Protein abundance is expressed as NSAF log2 fold-change on a color scale from blue to red where blue indicates the lowest and red the highest value, respectively. (B) Volcano plot of proteins identified in the mass spectrometry in ZM-treated vs CGS-treated CD8+ T-cells. The X axis indicates fold change (log2) and Y axis the p-value (-log10). The dotted lines indicates the cut-off separating proteins with p<0.05 (blue and red dots) from non-significant ones (black dots). Proteins that are interacting the least with Notch1 are shifted to the left of the plot whereas the ones interacting the most are shifted to the right. The bottom graph show the interaction between Notch1 and STS-1 (protein abundance) from the mass spectrometry analysis. (C) Primary isolated CD8+ T-cells were stimulated with anti-CD3/CD28 antibodies and treated with vehicle (control, DMSO) or 1µM CGS for 48h. Cbl-b was immunoprecipitated from protein lysates to detect its Tyr-phosphorylation. From top to bottom the panels show: Tyr-phosphorylation of Cbl-b, Cbl-b from IP samples, and Cbl-b, GAPDH from input samples. Densitometric (O.D.) results for Tyr-phosphorylate Cbl-b normalized by IP:Cbl-b are shown below the panel. Negative control refers to samples immunoprecipitated using beads but not antibody. (D) N1FL and N1TM were detected in protein lysates from primary CD8+ T- cells isolated from C57BL6 (WT) or STS-1/2 -/- (STS-1/2 KO) mice, activated with anti-CD3/CD28 antibodies and treated with vehicle (control, DMSO) or 1µM CGS for 72h. From top to bottom the panels show: N1FL, N1TM, STS-1 and β-actin. The graphs show averages ± standard deviation from three independent experiments. **p<0.01, two tailed T-test with equal variance.

### Genetic KO and pharmacologic inhibition of Cbl-b rescue Notch1 and T-cell functions from A2AR-mediated immunosuppression

Our data place Cbl-b at the core of an immunosuppressive pathway connecting A2AR, Notch1 and T-cell functions. Therefore, we asked if genetic KO or pharmacological inhibition of Cbl-b could be a strategy to increase T-cell functions and decrease immunosuppression by promoting Notch1. We treated CD8+ T-cells from Cbl-b KO mice with CGS and detected Notch1 level and INF-gamma (Fig. 5A,B). In agreement with our hypothesis, we found that CGS did not reduced Notch1 and INF-gamma in Cbl-b KO CD8+ T-cells (Fig. 5A,B), confirming that lack of Cbl-b prevents Notch1 downregulation and, in turn, promotes T-cell function. Considering these results, we asked if pharmacologic inhibition of Cbl-b could recapitulate what observed in Cbl-b KO CD8+ T-cells. To accomplish this, we tested the effect of novel small molecule compounds, designed to inhibit Cbl-b (supplementary fig. 3), in primary CD8+ T-cells. Consistent with our data in Cbl-b KO cells, we found that Cbl-b inhibitors, NTX-512, NTX-447 and NTX-307, rescue both full length and cleaved forms of Notch1 from CGS-mediated downregulation, suggesting Cbl-b inhibition prevents Notch1 downregulation (Fig. 5C). The effect of Cbl-b inhibitors on Notch1 resulted in rescue of proliferation, INF-gamma and Granzyme B (GNZB) production from CGS-mediated suppression and increased proliferation and production of cytokines compared to vehicle-treated CD8+ T-cells (Fig. 5D,E). The compounds showed dose-dependent effect on proliferation and cytokines production and remarkable potency as indicated by low EC50s both as single agents and against CGS (supplementary fig. 4A,B). Importantly, the compounds did not increase the production of INF-gamma in unstimulated CD8+ T-cells, indicating that they only boost the function of antigen-stimulated CD8+ T-cells (supplementary fig. 4C). These results strongly indicate that Cbl-b inhibition boosts Notch1 and T-cell functions and render CD8+ T-cells resistant to A2AR-mediated immunosuppression.

**Fig. 5.**
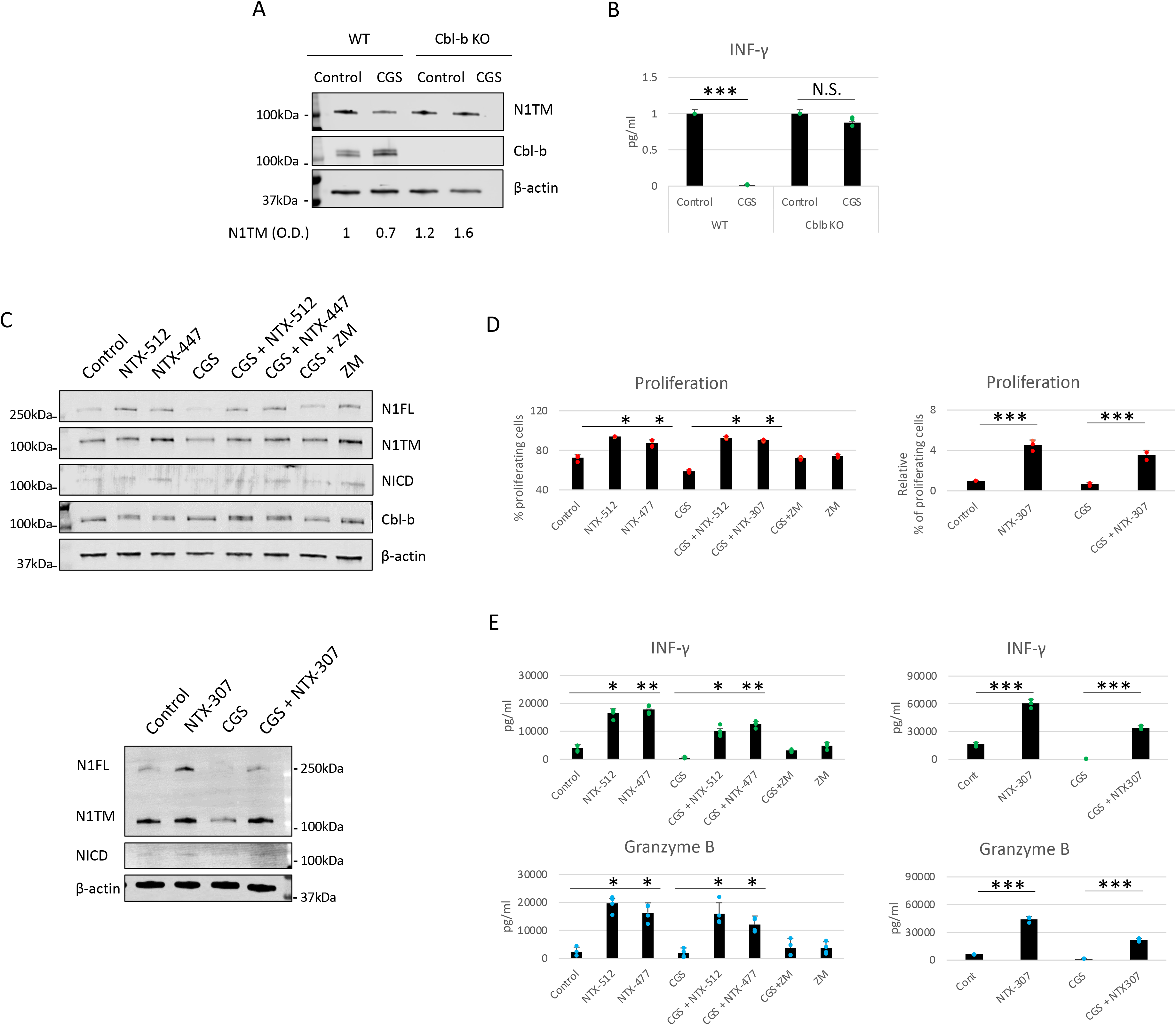
Genetic KO and pharmacologic inhibition of Cbl-b rescue Notch1 and T-cell functions from A2AR-mediated immunosuppression. (A) Primary CD8+ T- cells isolated from the spleen and lymph nodes of C57B16 (WT) or Cbl-b -/- (Cbl-b KO) mice were stimulated with anti-CD3/CD28 antibodies and treated for 72h with vehicle (control, DMSO) or NTX-512. Densitometric (O.D.) results for Notch transmembrane (N1TM) normalized by β-actin are shown below the panel. (B) Production of INF-γ and Granzyme B was analyzed using ELISA in supernatants from samples treated as described in (A). (C) Primary CD8+ T- cells isolated from the spleen and lymph nodes of C57B16 or FVB mice were stimulated with anti-CD3/CD28 antibodies and treated for 72h with vehicle (control, DMSO) or as indicated above the panels. All compounds were used at the concentration of 1µM. The top and bottom panels show Cbl-b inhibitors, NTX-512, NTX-447, and NTX-307, respectively. Protein lysates were analyzed for Notch1 full length (N1FL), Notch1 transmembrane (N1TM), Notch1 transcriptionally active form (NICD) and Cbl-b. (D) Proliferation was measure with flow cytometry by labelling primary CD8+ T-cells with CFSE in samples treated as in (C). (E) Production of INF-γ and Granzyme B was analyzed using ELISA in supernatants from samples treated as described in (C). The graphs show averages ± standard deviation from three independent experiments. *p<0.05, **p<0.01, ***p<0.001. two tailed T-test with equal variance. N.S. non-significant.

### Cbl-b inhibitors enhance Notch1-positive CD8+ T-cells anti-tumor responses

Since Cbl-b inhibition increases T-cell function and resistance to immunosuppression mediated by A2AR activation, we next wanted to test whether the effect of Cbl-b inhibition could translate in increased anti-tumor T-cells responses. For this purpose, we treated TNBC C0321 tumor-derived organoids with different concentrations of Cbl-b inhibitors and analyzed several readouts for anti-tumor activity. We found that Cbl-b inhibitors significantly induced cell death in organoids in a dose-dependent manner (Fig. 6A). This effect was absent in organoids derived from immunocompromised atymic Nu/Nu mice, suggesting that the anti-cancer activity of the compounds is immunological (Fig. 6A). Concomitantly, we found that Cbl-b inhibitors-treatment increased the production of INF-gamma in organoids cultures (Fig. 6B), a sign of increased CD8+ T-cell activation. Lastly, we labeled CD8+ T-cells and Notch1 in organoids and observed a significant infiltration of Notch1-positive CD8+ T-cells in organoids treated with Cbl-b inhibitors compare to control organoids (Fig. 6C). These Notch1-positive T-cells were surrounding cancer cells, possibly establishing immunological synapses for cancer cell killing. We also confirmed that Cbl-b inhibitors did not affect Notch1 in cancer cell lines, including C0321 (supplementary fig. 5). Overall, our results indicate that Cbl-b inhibition enhances CD8+ T-cell anti-cancer responses via Notch1 and show potential as single-agent cancer immunotherapy.

**Fig. 6.**
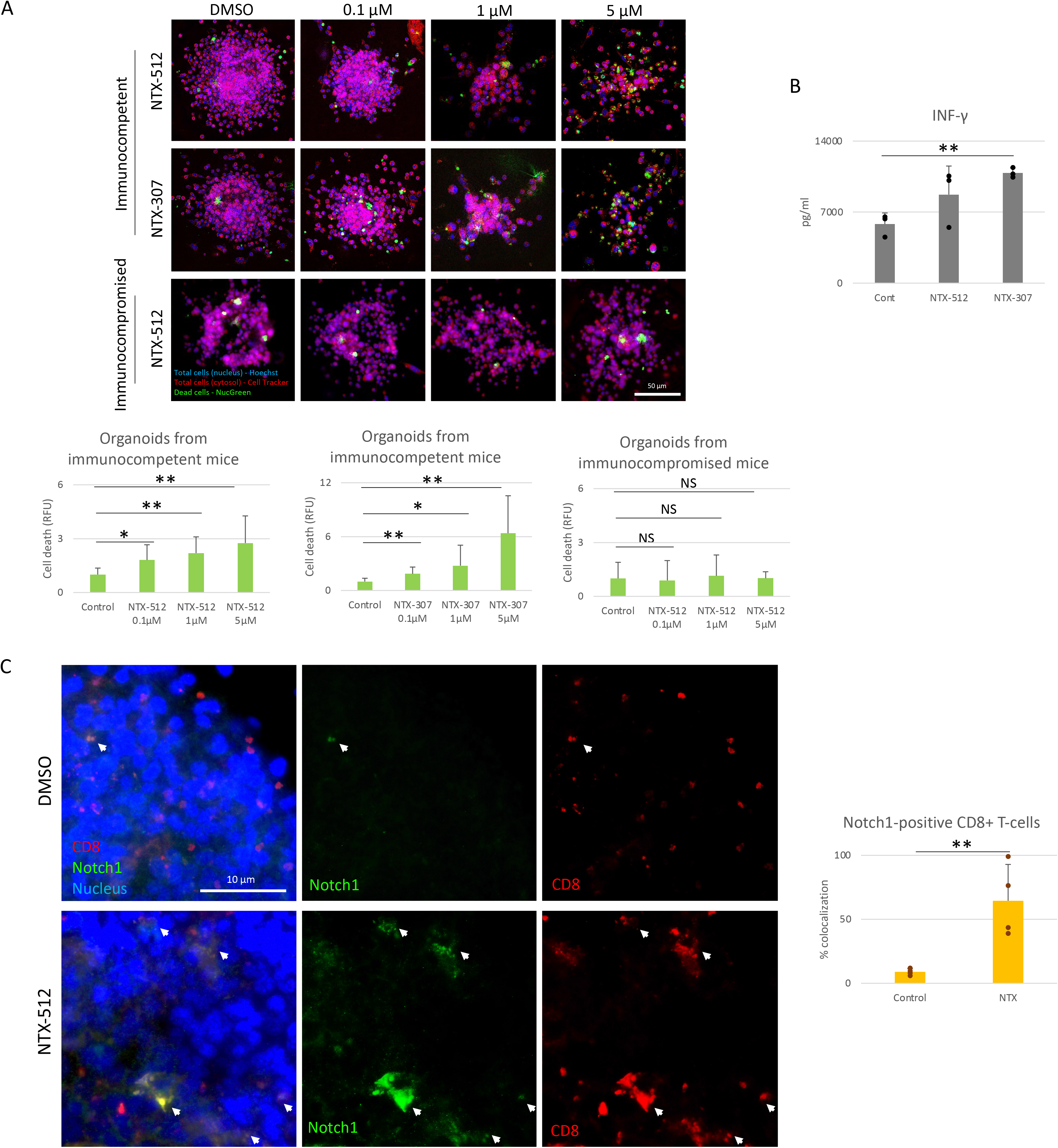
Cbl-b inhibitors enhance Notch1-positive CD8+ T-cells anti-tumor responses. (A) Organoids were obtained from a syngeneic TNBC mouse model, C0321, from immunocompetent FVB mice or immunocompromised athymic nude mice, cultured and treated for 4 days with vehicle (control, DMSO) or 0.1, 1 and 5µM of NTX-512 or NTX-307. On day 4, organoids were stained with Hoechst 33342 (nucleus), CellTracker Red (cytosol), NucGreen Dead 488 (dead cells) and imaged. Cell death was measured using ImageJ software. Production of INF-γ and Granzyme B was analyzed using ELISA in supernatants from organoids treated for 4 days with vehicle (control, DMSO) or 5µM NTX-512 or 5µM NTX-307. (C) C0321 organoids treated for 4 days with vehicle (control, DMSO) or 5µM NTX-512, were fixed, permeabilized and stained for CD8+ T-cells and Notch1 with primary anti-CD8, anti-Notch1 and secondary antibodies, and DAPI (nucleus). The colocalization of Notch1 and CD8 was measured using ImageJ software. All images were acquired on a BZ-x800 microscope with a 20x (A) and 60x (C) objectives. Scale bars length is indicated above each bar (µm). The graphs show averages ± standard deviation from ≥10 organoids from independent experiments. *p<0.05, **p<0.01, two tailed T-test with equal variance. NS, non-significant.

### Cbl-b inhibitors enhance immune-checkpoint immunotherapy

Immune-checkpoint inhibitor therapy has been approved for the treatment of certain tumors, including TNBC, but only a limited number of patients benefit from it (Haslam, 2019). Therefore, we tested if Cbl-b inhibitors could enhance the efficacy of anti-PD1/PDL1 immunotherapy. We treated organoids derived from two pre-clinical models of TNBC, C0321 and M-WNT (Zhang, 2014; Zhang, 2016; Dunlab, 2012), and colon cancer, MC38, with Cbl-b inhibitors (Fig. 7). We choose TNBC and colon cancer models since both tumors are known to have an immunosuppressive microenvironment (Hossain, 2021; Arora, 2018). The compounds were effective in inducing cell death in organoids both as single agents and in combination with anti-PD1/anti-PDL1 (Fig. 7). Interestingly, a marked synergy was seen between Cbl-b inhibitors and anti-PDL1 in all models (Fig. 7), thus highlighting the possibility that these treatments could be used for combinatorial immunotherapy. Our results show that Cbl-b inhibitors, by inducing Notch1-dependent CD8+ T-cells responses, have a promising potential as single-agent and combinatorial immunotherapy with checkpoint inhibitors. Cbl-b inhibition represents a new immunotherapeutic strategy that could be exploited to sensitize tumors to anti-cancer immune responses and treat tumors that are refractory to immunotherapy.

**Fig. 7.**
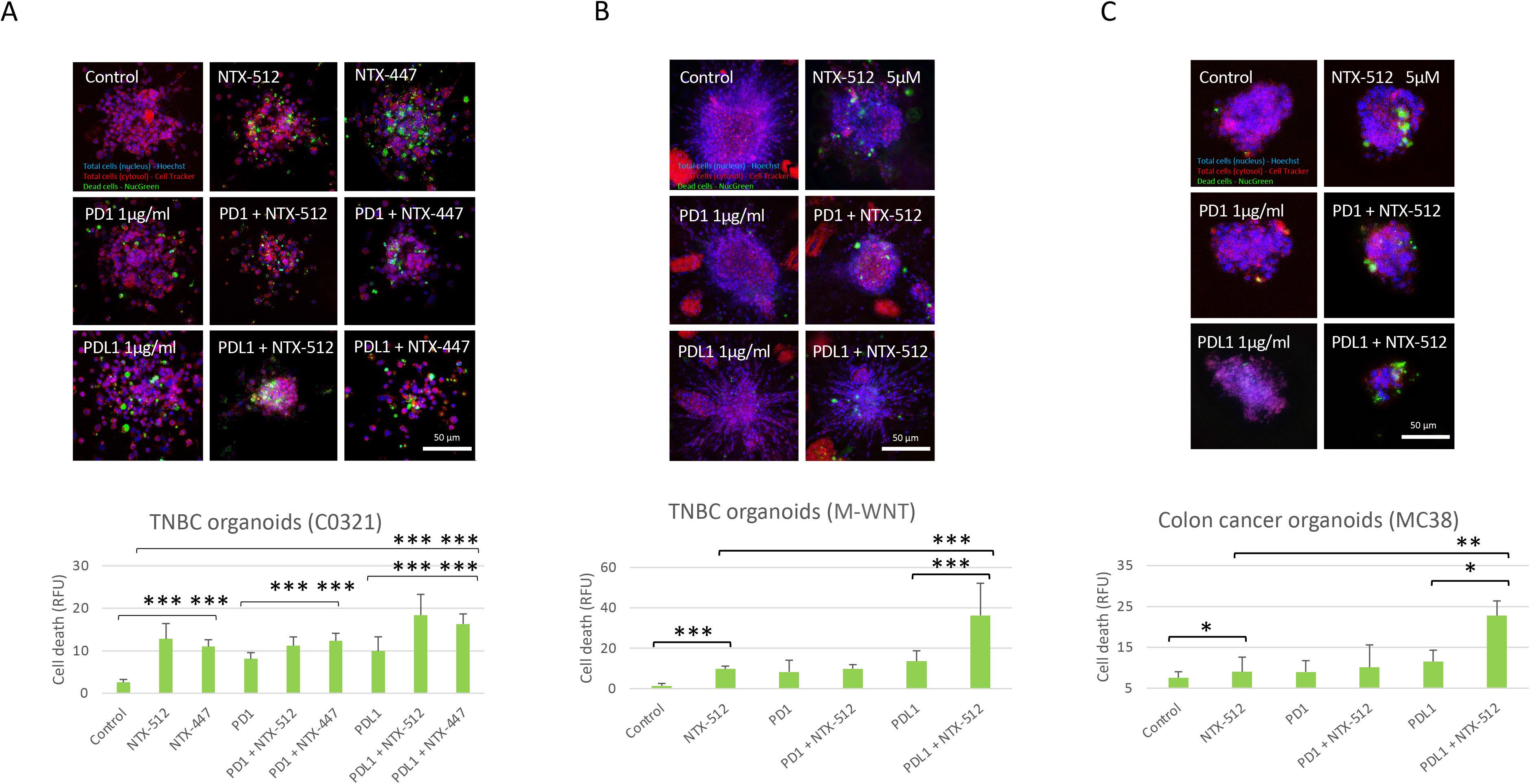
Cbl-b inhibitors enhance anti-PD1/PDL1 efficacy in tumor-derived organoids. Organoids were obtained from syngeneic pre-clinical TNBC, C0321 (A), M-WNT (B), and colon cancer, MC38 (C), models. Organoids were cultured in a collagen matrix and treated for 4 days with vehicle (control, DMSO) or 5µM of NTX-512 or NTX-477, alone or in combination with 1µg/ml anti-PD1 or 1µg/ml anti-PDL1. On day 4, organoids were stained with Hoechst 33342 (nucleus), CellTracker Red (cytosol), NucGreen Dead 488 (dead cells) and imaged. Cell death was measured using ImageJ software and all images were acquired on a BZ-x800 microscope with a 20x objective. Scale bars length is indicated above each bar (µm). The graphs show averages ± standard deviation from ≥10 organoids from independent experiments. *p<0.05, **p<0.01, ***p<0.001, two tailed T-test with equal variance. NS, non-significant.

## Discussion

Tumor-induced immunosuppression is a critical feature of cancer that allows evasion from the immune system (Hanahan, 2011). This is a major challenge for designing effective cancer immunotherapies that circumvent immunosuppression and are effective in a larger fraction of patients.

Our work describes a new regulatory pathway that is critical for immunosuppression in CD8+ T-cells and demonstrates that targeting this pathway is a promising strategy to overcome immunosuppression and enhance anti-cancer responses. We showed that activation of A2AR by adenosine, promotes Cbl-b-mediated Notch1 ubiquitination and degradation. STS-1 Tyr-phosphatase associates with Notch1 in response to A2AR activation and coordinates Cbl-b-mediated Notch1 degradation (Fig.8). Genetic KO of Cbl-b increases Notch1 levels. Pharmacological inhibition of Cbl-b results in increased Notch1, T-cell functions, anti-cancer response and resistance to immunosuppression in TNBC and colon cancer models. We also provided evidence illustrating that combinations of Cbl-b inhibition and immune-checkpoint inhibitors have enhanced efficacy in the same models. *In vivo* experiments were not attempted as the pharmacokinetics of the novel inhibitors we used is still under investigation. These chemical tools were selected based on specificity and *in vitro* activity.

**Fig. 8.**
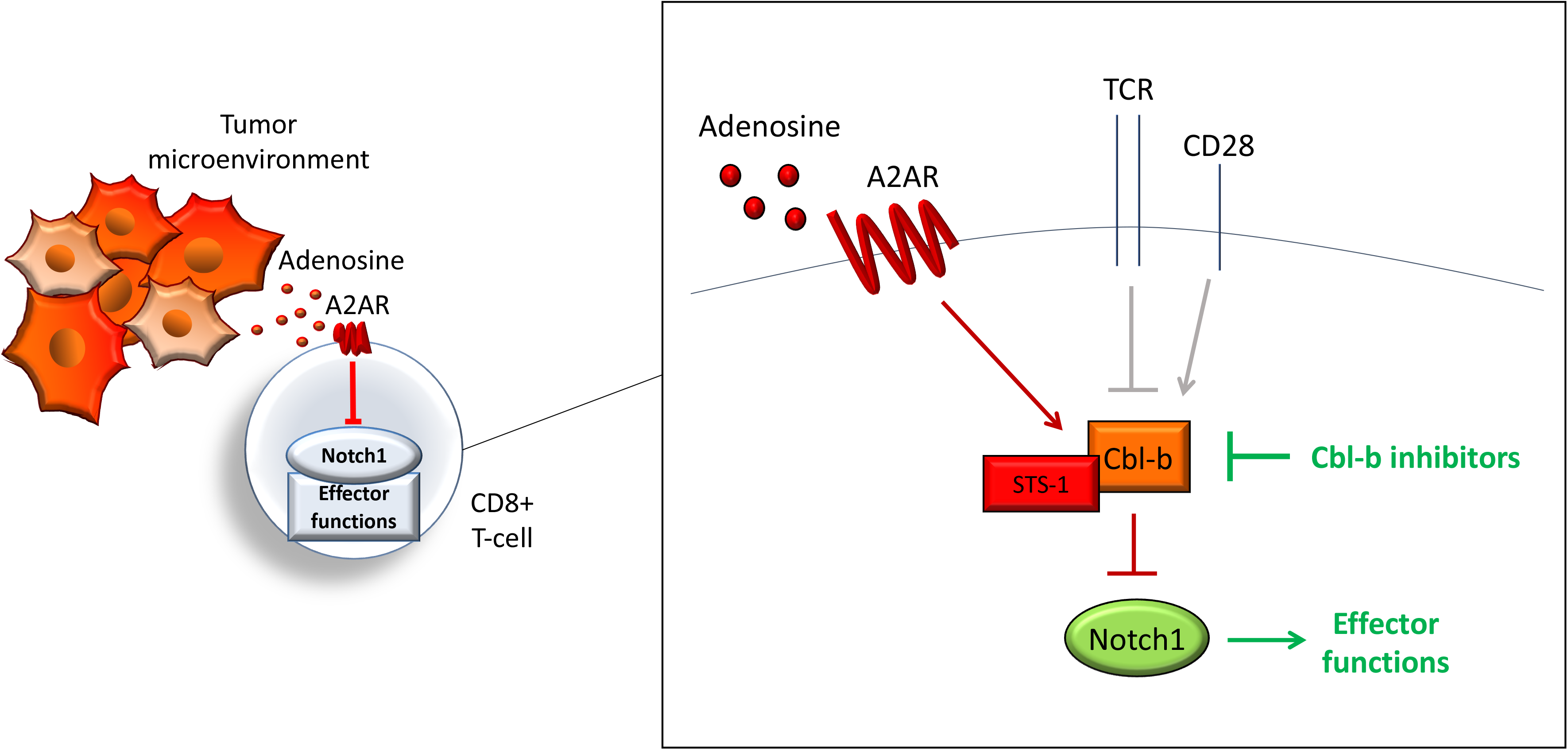
A2AR-Cbl-b-Notch1 pathway regulation. Adenosine induces immunosuppression in CD8+ T-cells in the tumor microenvironment by activating adenosine A2A receptor (A2AR), thus causing downregulation of Notch1 and suppression of effector functions. A2AR controls Notch1 levels by regulating the ubiquitin ligase Casitas B-lineage lymphoma b (Cbl-b), which mediates the ubiquitination and degradation of Notch1. A2AR promotes Cbl-b via Suppressor of T-cell receptor signaling 1 (STS-1) Tyr-phosphatase, possibly through Tyr-dephosphorylation of Cbl-b. A2AR, together with other receptors, including T-cell receptor (TCR) and co-stimulation receptor CD28, may control Notch1 levels in T-cells and, in turn, effector functions by modulating Cbl-b. This model places Cbl-b at the core of a regulatory axis, which, by integrating positive and negative signals from different receptors, determine the fate of Notch1 and effector functions. Pharmacological inhibition of Cbl-b blocks this axis and reactivates Notch1 and effector functions.

By describing A2AR-Cbl-b-Notch1 pathway in CD8+ T-cells, we showed for the first time a direct link between Cbl-b and adenosine-mediated immunosuppression, and placed Cbl-b and STS-1 at the center of Notch1 regulation in CD8+ T-cells. It is possible that Cbl-b mediates a constitutive degradative pathway that is switched on and off depending on whether the cell requires more or less Notch1. In T-cells Notch1 is activated in a ligand-independent manner through endocytosis (Steinbuck, 2018). Our work support a model in which the Cbl-b degradative pathway could be part of the ligand-independent endocytic pathway of Notch1 in T-cells, since endocytic regulation of Notch can either lead to activation or degradation (Steinbuck and Winandy, 2018; Monticone, 2021; Baron, 2012). In this endocytic pathway, Notch1 may be directed to degradation by Cbl-b-mediated ubiquitination, whereas other signals may instead direct Notch1 to activation, as observed in other systems (Shimizu, 2014; Wilkin, 2004). This pathway of degradation may be controlled in response to extracellular stimuli, including TCR activation and A2AR signaling, which both are known to regulate the level of Notch1 in T-cells (Sorrentino, 2019; Steinbuck, 2018; Cho, 2009; Palaga, 2003). For example, upon T-cell activation, signaling downstream of TCR may inhibit Cbl-b to elevate Notch1 and effector functions. Later, to avoid over-activation and exhaustion of T-cells, co-stimulatory signals (e.g. CD28) or immunosuppressive ones (e.g. A2AR) may promotes Cbl-b-mediated degradation of Notch1 and downregulation of effector functions (Fig.8). This would explain at least in part how Cbl-b controls the threshold for T-cell activation (Paolino, 2011; Chiang, 2000).

Previous work and our data, suggest that Cbl-b-mediated degradation may be regulated through phosphorylation. Cbl-b is negatively regulated by Tyr-phosphorylation by SHP-1 or Ser/Thr-phosphorylation by PKC-theta in response to CD28 co-stimulation, but dephosphorylated and promoted in response to TCR stimulation (Xiao, 2015; Gruber, 2009; Zhang, 2002). Accordingly, we found evidence that Cbl-b was dephosphorylated by STS-1, and promoted upon A2AR activation. It is very likely that the phosphorylation of different residues or a different phosphorylation status of Cbl-b may increase or decrease Cbl-b function. For instance, Tyr-phosphorylation of specific residues of Cbl-family of ubiquitin ligases was instead found to increase their activity (Kassenbrock, 2004). Therefore, it would be interesting to identify the residues that are dephosphorylated by STS-1. It is also possible that STS-1 and other phosphatases/kinases in this network regulate the phosphorylation of Notch1, as phosphorylation can also act as a signal for target degradation or activation (Morrugares, 2020; Xiao, 2015). If Cbl-b is controlled by phosphorylation, it is also possible that a Tyr-kinase antagonizes STS-1 function, by phosphorylating and inhibiting Cbl-b. Lck Tyr-phosphorylates Cbl-b in T-cells (Xiao, 2015). Lck is inhibited by cAMP level via PKA and Csk (Vang, 2001). It is possible that activation of A2AR, which increases the level of cAMP, inhibits Lck-mediated phosphorylation of Cbl-b (supplementary fig. 6). This is in line with the observation that Lck interacts with Notch1 and Lck inhibition in T-cells completely blocks Notch1 activation (Steinbuck, 2018; Sade, 2004), possibly by promoting Cbl-b-mediated degradation of Notch1. Overall, our work supports the idea that antagonistic phosphorylating signals from TCR and other receptors, like A2AR, control Cbl-b and in turn Notch1-effector functions, thus regulating the threshold of T-cell activation. This network that integrates extracellular stimuli with intracellular signals via phosphorylation and ubiquitination may participate in other regulatory pathways in T-cell activation.

Despite the successful application of immunotherapy, response rates remain limited (Haslam, 2019). They could be increased by new immunotherapies refractory to different forms of immunosuppression and that can work as single-agent or in combination with existing immunotherapies. Our work, together with previous studies (Sorrentino, 2019; Sierra, 2014), supports the idea that therapies that reactivate Notch1 in T-cells could be used to tune T-cells against immunosuppression and enhance anti-cancer immune responses (Fig. 9). Specifically, we described a new pathway that is amenable to drug targeting and has promising selectivity for Notch in T-cells versus cancer cells, a feature that is very important for Notch-targeted therapies (Majumder, 2021). We observed increased/restored effector functions when Notch1 is reactivated in T-cells. We also observed that reactivation of Notch1 primes T-cells to attack cancer cells in tumor-derived organoids. By reactivating Notch1 in T-cells we aim to lower the threshold of T-cell activation to make suppressed T-cells more reactive to tumor recognition when infiltrating in the tumor microenvironment (Fig. 9). It remains to be determined whether this strategy can work both locally in the tumor microenvironment and systemically, as it is becoming clear that systemic immunosuppression is critical in cancer patients (Hiam-Galvez, 2021). Our results in organoids from TNBC and colon cancer, two tumor types which can suppress immune responses (Hossain 2021; Arora, 2018), suggest that Notch1 reactivation and Cbl-b inhibition could also be a viable strategy to sensitize “cold” tumors to cancer immunotherapy. Another promising application of this immunotherapeutic strategy, could be adoptive T-cell therapy. Resistance to immunosuppression plays a critical role in these therapies and both Notch1 expression and Cblb deletion were found to enhanced the efficacy of adoptive T-cell transfer and CAR-T cell therapy (Kumar, 2021; Sierra, 2014). Our work presents a new class of candidate immunotherapeutic compounds, Cbl-b inhibitors, that enhance anti-cancer responses and resistance to immunosuppression. We show that Cbl-b inhibitors are effective as single-agents or in combination with anti-PDL1/PD1 in organoids derived from pre-clinical cancer models. Cbl-b inhibitors are currently in the discovery phase of development. If their efficacy will be confirmed *in vivo* in pre-clinical mouse models, these compounds certainly hold a promising future development as a new generation of cancer immunotherapeutics.

**Fig. 9.**
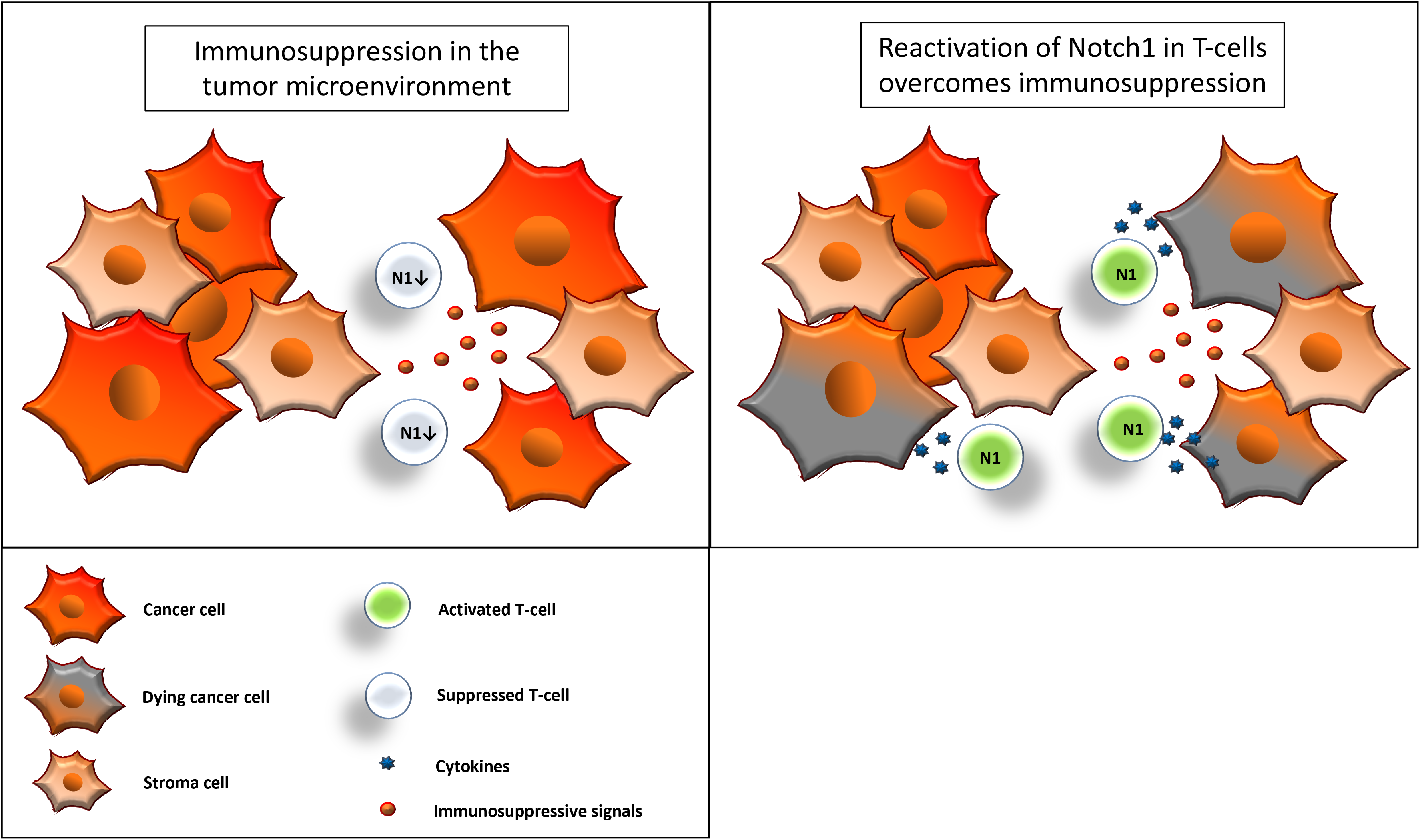
Reactivation of Notch1 in T-cells overcomes immunosuppression. Immunosuppressive signals induce suppression in CD8+ T-cells in the tumor microenvironment. Some of these signals, including adenosine, may suppress CD8+ T-cells effector functions by downregulation of Notch1. Reactivation of Notch1, with Cbl-b inhibitors or other strategies, may tune CD8+ T-cells against immunosuppression and enhance effector functions, ultimately promoting anti-cancer responses.

Our work described for the first time a critical immunosuppressive pathway between A2AR, Cbl-b and Notch1. We showed that promoting Notch1 signaling by blocking Cbl-b-mediated degradation results in a robust increase in anti-cancer T-cell responses and resistance to immunosuppression. Our findings demonstrate that targeting Cbl-b-Notch1 axis represents a promising novel immunotherapeutic strategy to boost anti-cancer T-cell responses.

## Materials and Methods

### Mice

C57BL/6 and FVB mice (6-8 week-old) were purchased from The Jackson Laboratory (Bar Harbor, ME) and housed in specific pathogen-free conditions in a 12h light/12h dark cycle with food and water available ad libitum. For isolation of CD8+ T-cells, spleens and lymph nodes were aseptically harvested post-euthanasia. For the orthotopic TNBC models 2×10^6 million C0321 or M-WNT cells were injected into the 4th mammary fat pad of C57BL/6 or FVB mice, respectively. For the syngeneic colon adenocarcinoma model, 2.5×10^5 MC-38 were injected subcutaneously in C57BL/6 mice. All experiments involving mice were approved by the Institutional Animal Care and Use Committee (IACUC) at LSUHSC (New Orleans, LA).

### Cell lines

C0321 and M-WNT TNBC cell lines were developed as described in Zhang, 2014, Zhang, 2015 and Dunlab, 2012. MC-38 colon adenocarcinoma cell line and 293T cells were purchased from ATCC (Manassas, VA). C0321 and 293T cells were cultured in DMEM and M-WNT and MC-38 cells in RPMI. Both media were supplemented with 10% fetal bovine serum, 4 mM L-Glutamine, 50 U/ml penicillin and 50 µg streptomycin (Gibco).

### Primary CD8+ T-cells

CD8+ T-cells were aseptically isolated from the spleen and lymph nodes of C57BL/6 or FVB (6-8 weeks old) mice using the negative selection Easysep mouse CD8+ T-cell isolation kit (StemCell Technologies) according to the manufacturer’s instructions. Spleens and lymph nodes from Cbl-b KO (*Cbl-b ^-/-^*) were kindly provided by Dr. J. Chiang (NCI-NIH; Chiang, 2000) and STS-1/STS-2 double KO (*STS-1 ^-/-^ / STS-2 ^-/-^*) by Dr. N. Carpino (Stony Brook University; Carpino, 2004). CD8+ T-cells were cultured in RPMI supplemented with 10% fetal bovine serum, 4 mM L-Glutamine, 50 U/ml penicillin, 50 µg streptomycin and 50 µM 2-mercaptoethanol. The cells were activated in plates coated with anti-mouse CD3ε and anti-mouse CD28 antibodies (1 µg/ml, BD Biosciences) for up to 72h.

### Tumor-derived organoids and imaging

Organoids were derived from C0321, M-WNT or MC-38 tumors in FVB and C57BL/6 mice. To establish organoids cultures, tumors were minced and digested at 37°C in FBS-free DMEM/F12 Glutamax containing 1mg/ml type IV collagenase (Gibco). Digested tumors were passed through a 100 μm and a 70 μm strainer to isolate organoids of 70-100 μm in size. The organoids were resuspended in type I rat tail collagen gel (Gibco) and plated in 8-well chambered coverslips (µ-slide 8 well, Ibidi). The organoids cultures were hydrated with DMEM/F12 Glutamax supplemented with 5% FBS, 50 U/ml penicillin and 50 µg streptomycin. Treatments were added directly into the medium after plating the organoids-collagen cultures. Cells in organoids were stained using CellTracker™ Red CMTPX (cytosol, Invitrogen) and Hoechst 33342 (nucleus, BD Biosciences), and dead cells were labelled using a cell membrane impermeable nucleic acid dye, NucGreen™ Dead 488 ReadyProbes™ (Invitrogen). Infiltrating CD8+ T-cells were stained in organoids using rat anti-mouse CD8α PE-conjugated antibody (Clone 53-6.7, BD biosciences). For CD8-Notch1 co-localization experiments, organoids were fixed in 2% formalin, permeabilized in 0.2% Triton and block with 2.5% goat serum and Fc block (BD Biosciences). CD8+ T-cells were stained with rat anti-mouse CD8α PE-conjugated antibody (Clone 53-6.7, BD biosciences) and Notch1 with rabbit anti-Notch1 (D6F11, Cell Signaling Technology) primary antibody, and goat anti-rat Texas Red (Invitrogen), goat anti-rabbit Alexa Fluor 488 (Invitrogen) secondary antibodies, respectively. Cell nuclei were stained using DAPI.

Organoids were imaged at day 0-6 using a BZ-x800 microscope (Keyence) with 4x, 20x or 60x objectives. To measure the size of organoids over time, bright field images of a given organoid were taken at day 0, 4 and 6 of culture using the multi-point tool of BZ-x800, which allows to save a specific position in the culture plate. The area of the organoids was measured using ImageJ. To quantify cell death in organoids, the area positive for dead cell staining was measured and normalized by the total area of organoids using BZ-x800 analyzer software; or the fluorescence intensity of the dead cell staining was measured and normalized by the area of the organoids using ImageJ. The same method was used to quantify the infiltration of CD8+ T-cells, by measuring the CD8+ area/ fluorescence intensity and normalizing by the total organoid area. To quantify the co-localization between Notch1 and CD8, Notch1 fluorescence intensity was measured within the CD8+ areas of at least three different images per sample using ImageJ. Full co-localization fluorescence intensity was set to a value of 100 and the fluorescence intensity values obtained were expressed in percentages accordingly.

### Compounds and potency studies

A2AR agonist, CGS-21680, and antagonist, ZM-241385, were purchased from Tocris. Cbl-b inhibitors, NTX-512, NTX-447 and NTX-307, were produced by Nimbus Therapeutics. NTX compounds were designed using a structure-guided approach to inhibit Cbl-b enzymatic activity. This approach ensures exquisite selectivity towards Cbl-b and c-Cbl, another member of the Cbl-family of ubiquitin ligases. The potency of NTX compounds to inhibit Cbl-b was assessed by time-resolved measurement of fluorescence with fluorescence resonance energy transfer technology (TR-FRET). Briefly, recombinant human Cbl-b (aa 36-427) was expressed in *E. coli*, purified and biotinylated *in vitro*. Recombinant human Src (aa 254-536)-Zap-70 (aa 281-297) fusion protein was expressed in *E. coli* and purified. Recombinant human UBE2D2(C85K) was expressed in *E. coli*, purified, ubiquitinated and BODIPY labelled *in vitro*. The compounds were dissolved in DMSO (typically at 10-20mM) and a ten-point half log dilution series was prepared using acoustic dispensing. The assay was performed by adding Cbl-b enzyme and Src-Zap/ATP (enzyme reaction) in the presence of UBE2D2(C85K)-Ub-FL-BODIPY, Streptavidin-Tb (binding reaction). The assay signal was measured at 520nm on an Envision plate reader, with reference signal at 620nm. Data was normalized using high and low assay controls: % Inhibition =100-(100*((high control) - unknown) / (high control - low control)). A 4-parameter dose-response equation was used to fit the normalized dose-response data and derive an IC50 for the compounds.

### Constructs and transfection

The constructs used were as follows: all HA-tagged Cbl-b constructs were a gift from Dr. Stanley Lipkowitz (Ettenberg, 2001; Ettenberg, 1999) while the His-tagged ubiquitin construct was from Dr. Dirk Bohman (Treier, 1994). NTM and all Myc-tagged Notch1 deletion constructs have been previously described (Jeffries, 2000; Aster, 1997). Lipofectamine 3000 (Invitrogen) was used to transfect 293T cells following the manufacturer’s instructions.

### Luciferase reporter assay

Luciferase assays were performed using the Dual Luciferase assay kit (Promega). 293T cells were transfected with Notch and Cbl-b constructs as indicated. All cells received a Hes-luciferase construct and a Renilla luciferase expression vector (pRL-CMV). Cells were cultured for 48 hours, harvested and lysates were made as per the manufacturer’s protocol. Reporter gene transcription was measured using a luminometer.

### Immunoprecipitation and western blot

Primary CD8+ T-cells were lysed in RIPA lysis buffer (Santa Cruz Biotechnology) for western blot or in Pierce IP lysis buffer (Thermo Fisher) for IP, supplemented with 1 mM Protease Inhibitor Cocktail (Thermo Fisher), 1 mM PMSF and 1 mM sodium orthovanadate. For IP, lysates were pre-cleared using Pierce Protein A/G Magnetic beads (Thermo Fisher) for 1 hour. Pre-cleared lysates were incubated over night at 4°C with end-over-end rotation with 2 µg of anti-Notch1 (D1E11, Cell Signaling) or anti-Cbl-b (G1, Santa Cruz Biotechnology) or rabbit IgG control antibody (Thermo Fisher). Pierce Protein A/G Magnetic beads (Thermo Fisher) were added and incubated for 1 hour at room temperature with end-over-end rotation. The beads were washed three times with lysis buffer and two times with deionized water at RT, resuspended in 2X Laemmli sample buffer (Biorad) and incubated at 95°C for 5 minutes to elute the immunoprecipitated proteins. For mass spectrometry, IP samples were prepared using Pierce MS-compatible Magnetic IP kit (Thermo Fisher) according to the manufacturer’s instructions.

293T cells were transfected with constructs as indicated, cultured for 48 hours and lysates were prepared using a lysis buffer containing 50 mM Hepes (pH 7.8), 1% NP-40, 250 mM NaCl, Pefabloc (0.125 mM, Fluka) and approximately 1 mg of protein was used per IP. For the ubiquitination assay, the proteosomal inhibitor lactacystin (20 μM, Kamiya Biomedical Corp.) was added to the culture media for 8 hours before preparing protein lysates with the above lysis buffer supplemented with N-Ethylmaleimide (25mM, Sigma). Lysates were pre-cleared using Protein G sepharose beads (GE Healthcare) for 1 hour at 4°C and, depending upon the experiment, 4-6 µg of anti-Notch1 (C-20, Santa Cruz Biotechnology), anti-HA tag (Cell Signaling) or rabbit IgG control antibody (Santa Cruz Biotechnology) was added to pre-cleared lysates. After incubating on ice for 2 hours, 30 µl of Protein G sepharose beads were added per tube and incubated for 1 hour at 4°C with end-over-end rotation to pull down antigen-antibody complexes. Beads were washed thoroughly with lysis buffer at RT, resuspended in 2X Laemmli sample buffer (Biorad) and incubated at 95°C for 5 minutes to elute the immunoprecipitated proteins.

For western blotting, protein lysates were resolved on 4-15% or 7.5% SDS-PAGE gels (Biorad) and transferred on to PVDF membranes (Millipore). Blots were incubated overnight with primary antibody diluted in Intercept blocking buffer (Licor). The next day, blots were incubated with an appropriate HRP-conjugated secondary antibody for chemiluminescence detection, or with IRDye fluorescent secondary antibodies (Licor) for fluorescence detection, for 1 hour at RT. Proteins were visualized by developing the blots with ECL reagent (Biorad) or imaged on an Odyssey scanner (Licor). The following primary antibodies were used for western blot in this study: anti-Notch1 (D1E11, Cell Signaling, or mN1A, Novus, or C-20, SCBT) for Notch1 full length and cleaved forms; anti-cleaved Notch1 Val1744 (D3B8, Cell Signaling or PA5-99448, Invitrogen-Thermofisher) for NICD; anti-Cbl-b (G1, Santa Cruz Biotechnology or 12781-1-AP, Proteintech); anti-STS-1 (19563-1-AP, Proteintech); anti-β-actin (AC-15, SCBT); anti-phospho-Tyr (PY99, STCB); anti-HA tag (Clone 6E2, Millipore).

### Mass spectrometry

Mass spectrometry analyses was performed by Dr. Samuel Mackintosh’s team as part of the IDeA National Resource for Quantitative Proteomics Voucher program. The samples were trypsin-digested and analyzed through data independent acquisition (DIA) quantitative proteomic platform in an Orbitrap Exploris 480 mass spectrometer.

### ELISA assay

Cytokine production was measured in supernatants of primary CD8+ T-cells activated and treated as explained above, using ELISA kits (Invitrogen) according to the manufacturer’s instructions.

### Proliferation assay

Proliferation was measured by labelling primary CD8+ T-cells with 1 μM carboxyfluorescein diacetate succinimidyl ester (CFSE, Thermo Fisher) for 10 min at 37°C before activation and treatments. CFSE fluorescence was measured by flow cytometric analysis on a Gallios cytometer (Beckman Coulter) and data analyzed using Kaluza software (Beckman Coulter).

### Statistics

Two-tailed unpaired Student’s *t*-test with Bonferroni correction was used for comparisons between groups. P values ≤ 0.05 were considered significant.

## Acknowledgments

This work was supported in part by P20CA233374. We thank Nimbus Therapeutics for providing Cbl-b inhibitors; Dr. Jeffrey Chiang for providing Cbl-b KO T-cells; Dr. Samuel Mackintosh, his team and the IDeA proteomics program for the mass spectrometry analysis; Dr. Stephen D. Hursting for providing M-WNT cells; Dr. Maira Di Tano, Dr. Cristina Melero, Dr. Francesca Carrieri, and Dr. Ramesh Thylur for providing insightful discussions; Dr. Augusto Ochoa, Dr. Kriss Reiss and their research groups for providing technical assistance.

## Conflict of interest

The authors declare that they have no conflict of interest.

---

**Supplementary Fig. 1.**
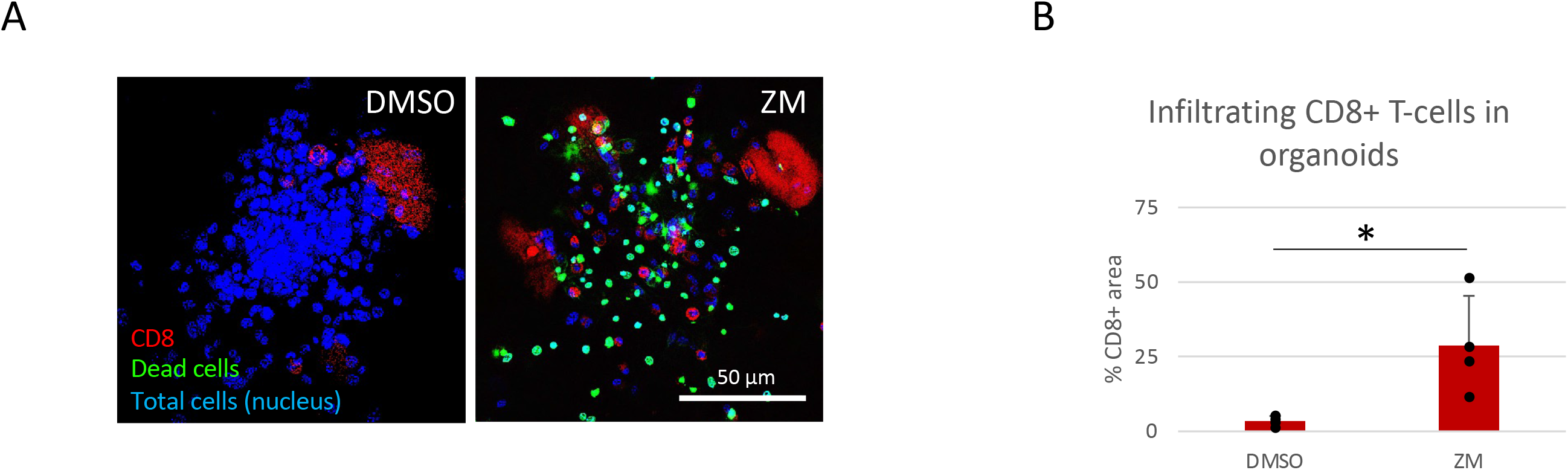
CD8+ T-cells infiltration in organoids. (A) C0321 organoids were stained for CD8+ T-cells using a fluorochrome-conjugated anti-CD8 antibody, Hoechst 33342 (nucleus) and NucGreen Dead 488 (dead cells), after 4 days in culture with vehicle or 5µM ZM. (B) The percentage of CD8+ T-cells was calculated using BZ-x800 analyzer software. Scale bars length is indicated above each bar (µm). The graphs show averages ± standard deviation from ≥4 independent experiments. *p<0.05, two tailed T-test with equal variance. NS, non-significant.

**Supplementary Fig. 2.**
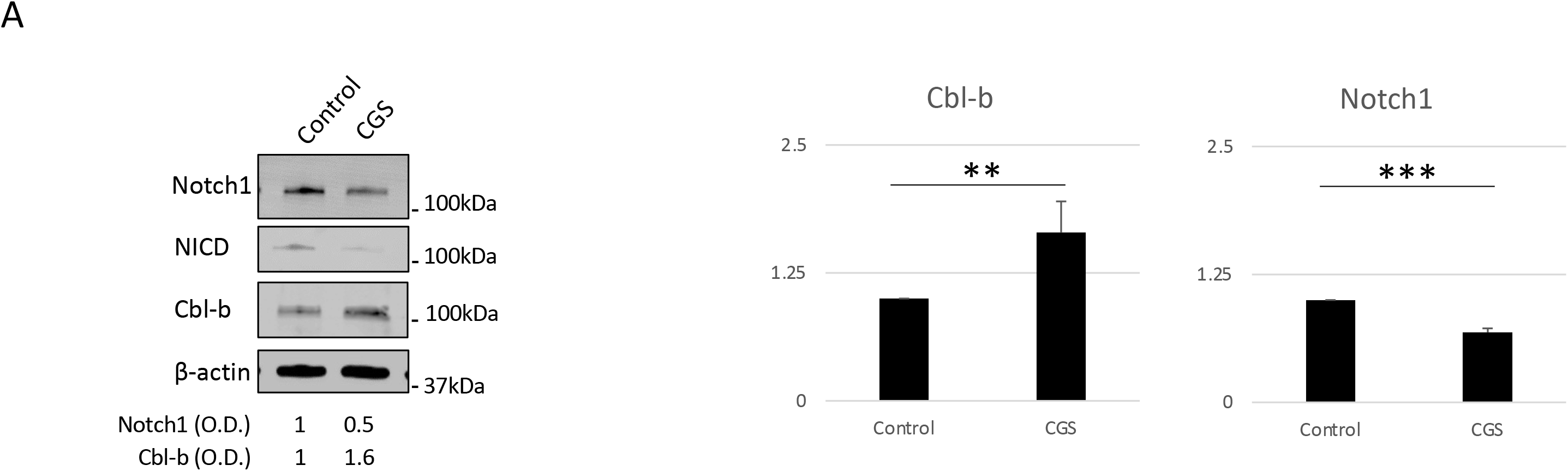
Cbl-b protein level is increased by A2AR activation in CD8+ T-cells. (A) Primary CD8+ T-cells isolated from the spleen and lymph nodes of C57B16 or FVB mice were stimulated with anti-CD3/CD28 antibodies and treated for 72h with vehicle (control, DMSO) or CGS 1µM. Protein lysates were analyzed for Notch1 transmembrane (N1TM), Notch1 transcriptionally active form (NICD) and Cbl-b. Densitometric (O.D.) results for Cbl-b normalized by β-actin are shown below the panel. (B) Densitometry analysis for Cbl-b protein levels. The graphs show averages ± standard deviation from three independent experiments. **p<0.01, ***p<0.001. two tailed T-test with equal variance.

**Supplementary Fig. 3.**
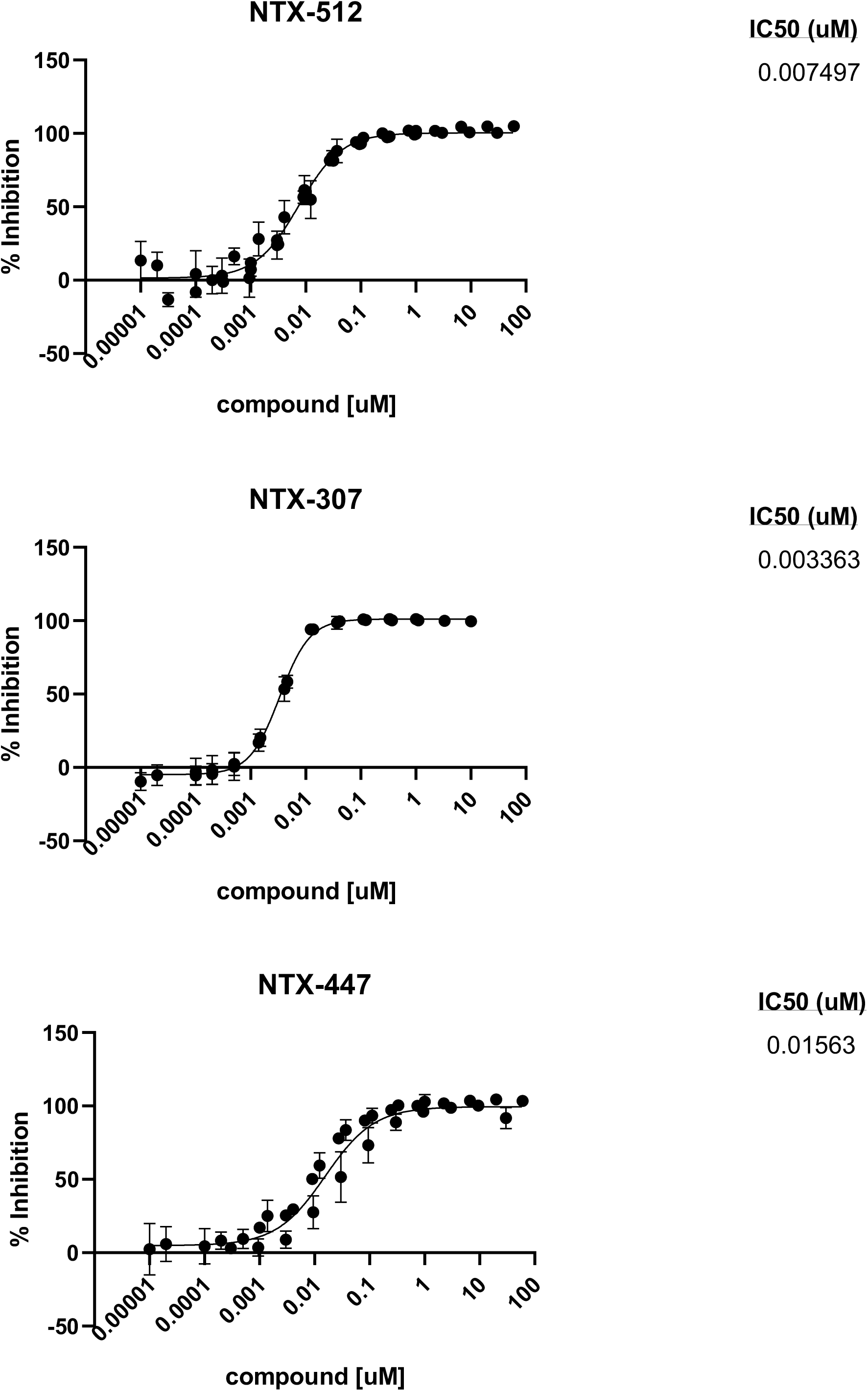
Potency studies of NTX compounds. The potency of NTX compounds to inhibit Cbl-b was assessed by time-resolved measurement of fluorescence with fluorescence resonance energy transfer technology (TR-FRET). Dose-response curves and IC50s were calculated using a 4-parameter dose-response equation. The X axis of the graphs show concentrations (µM) in log10. The Y axis show the responses (Cbl-b inhibition) expressed as percentages. Data was normalized using high and low assay controls: % Inhibition =100-(100*((high control) - unknown) / (high control - low control)).

**Supplementary Fig. 4.**
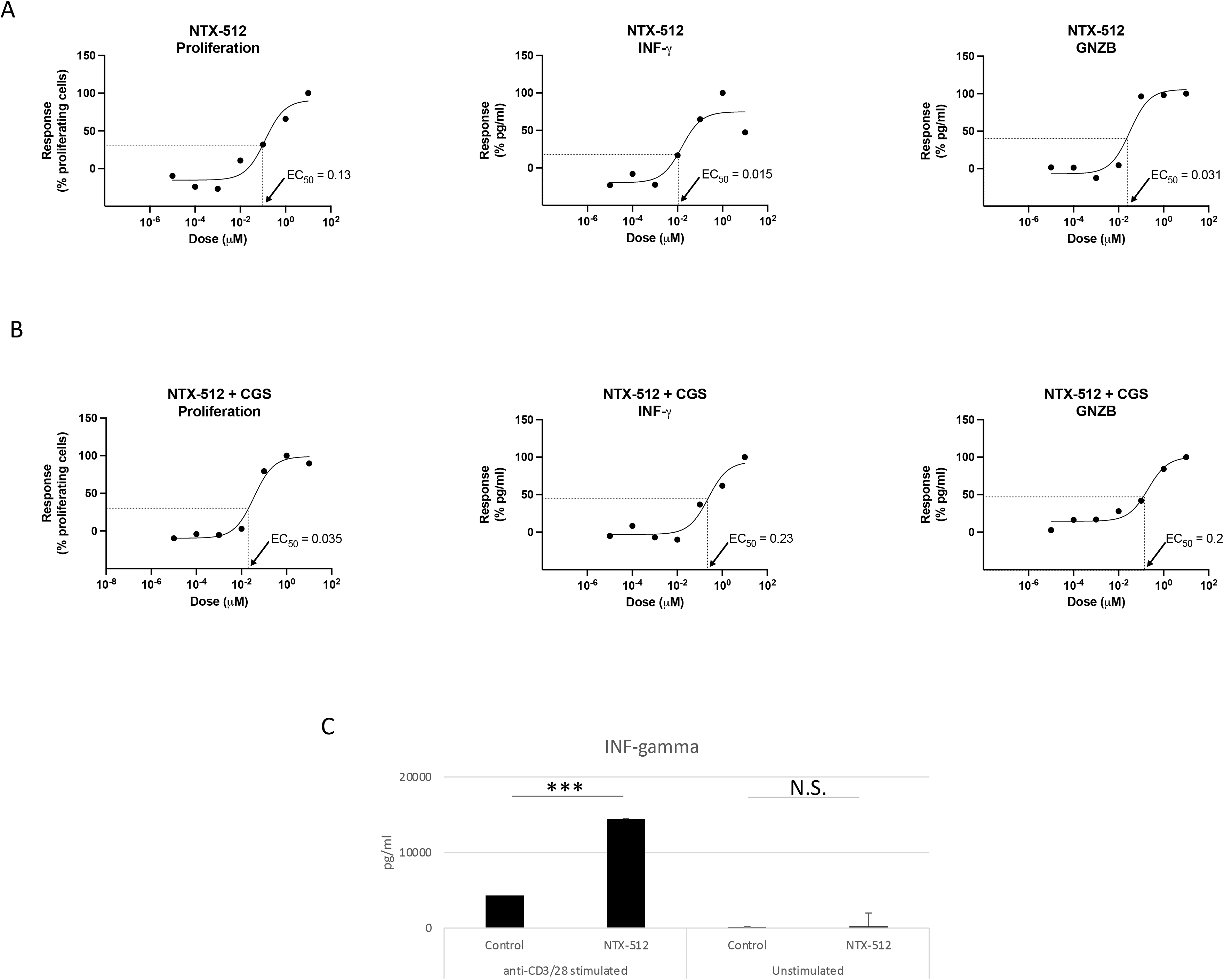
Cbl-b inhibitors show a dose-response effect in activated T-cells and no effect in unstimulated T-cells. Primary mouse CD8+ T-cells isolated from C57BL6 were stimulated with anti-CD3/CD28 and treated for 72h with different concentrations of NTX-512 alone (A) or in combination with 1µM CGS (B). Proliferation was measure with flow cytometry by labelling cells with CFSE and production of INF-γ and Granzyme B was analyzed using ELISA in supernatants. The graphs show averages ± standard deviation from three independent experiments. Dose-response curves and EC50s were calculated using Graphpad Prism software. The X axis of the graphs show concentrations (µM) in log10. The Y axis show the responses (proliferation or cytokine production) expressed as percentages. For clarity, the highest response value was set to 100% and the other values were set accordingly. (C) Production of INF-γ was analyzed using ELISA in supernatants from CD8+ T-cells stimulated with anti-CD3/CD28 or unstimulated and treated with 1µM NTX-512 or vehicle-treated (Control, DMSO). The graphs show averages ± standard deviation from three independent experiments. ***p<0.001. two tailed T-test with equal variance. N.S. non-significant.

**Supplementary Fig. 5.**
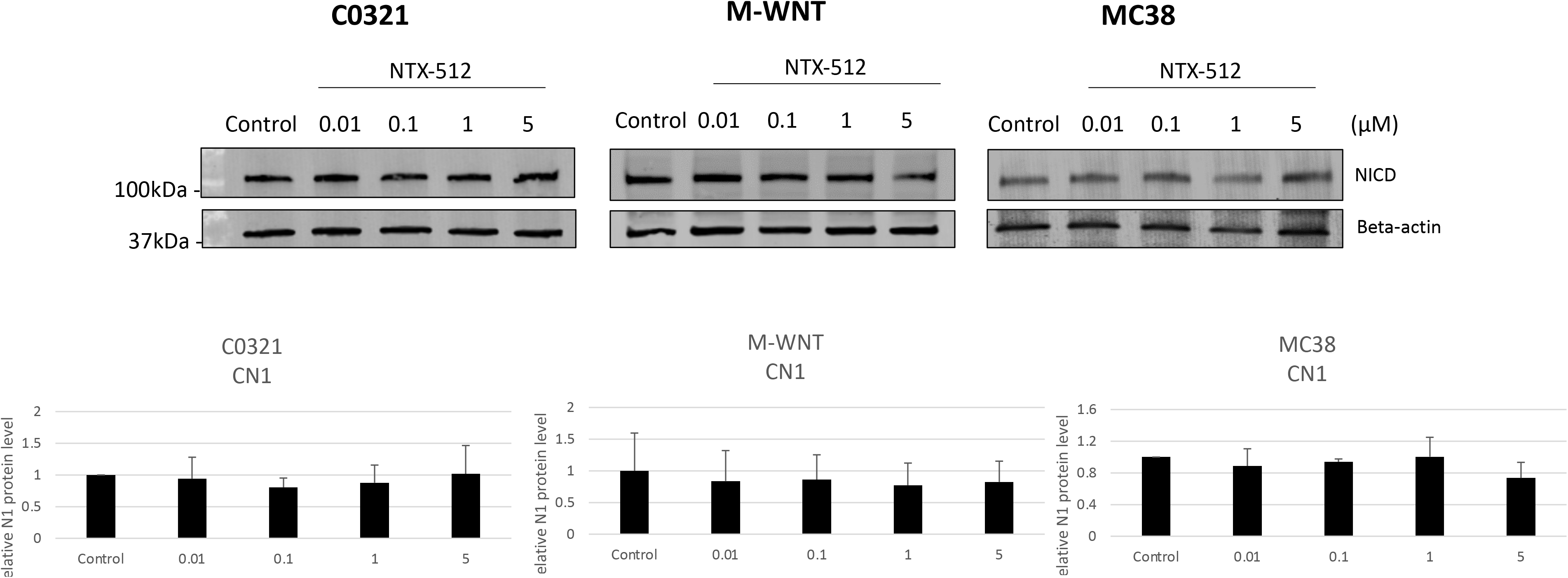
Cblb inhibitors do not modify Notch1 in cancer cells. TNBC cancer cell lines, C0321, M-WNT, and colon cancer, MC38, were cultured in complete medium and treated for 48h with vehicle (control, DMSO) or 0.01, 0.1, 1 and 5µM of NTX-512. Protein lysates were analyzed for Notch1 transcriptionally active form (NICD). The graphs show the densitometric analysis of NICD normalized by β-actin. The graphs show averages ± standard deviation from three independent experiments. Two tailed T-test with equal variance identify no significant differences in NICD in the samples.

**Supplementary Fig. 6:**
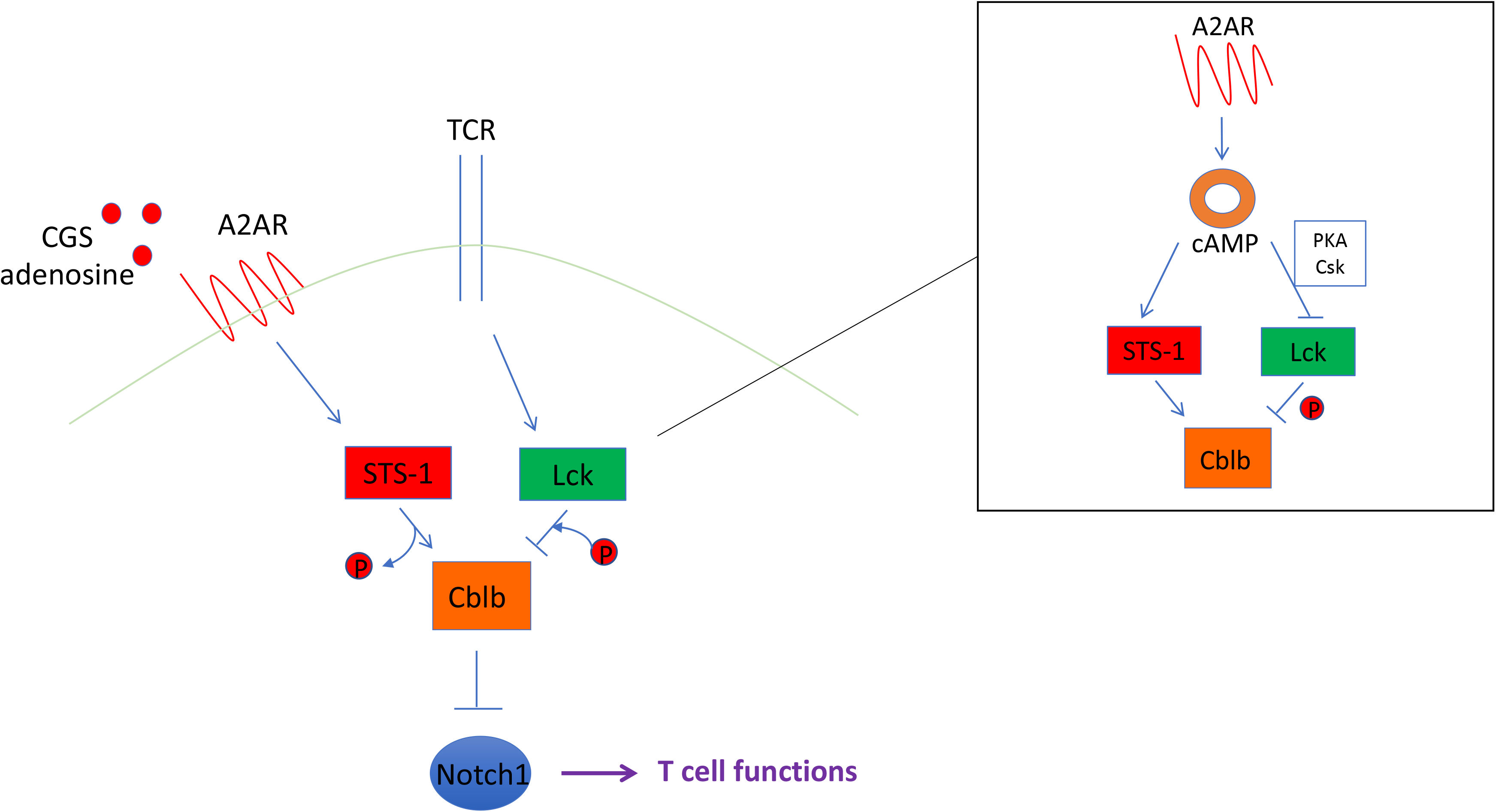
Working model of regulation of Cbl-b-Notch1 axis via phosphorylation. STS-1 may dephosphorylate and promote Cbl-b, whereas Lck Tyr-kinase may antagonize STS-1 function, phosphorylate and inhibit Cbl-b, thus preventing Notch1 degradation. Lck is inhibited by cAMP level via PKA and Csk. A2AR activation, which increases the level of cAMP, may inhibit Lck-mediated phosphorylation of Cbl-b.

